# Light and chloroplast redox state modulate the progression of tobacco leaf infection by *Pseudomonas syringae* pv *tabaci*

**DOI:** 10.1101/2025.01.06.631282

**Authors:** Rocío C. Arce, Mariana Demarchi, Nicolás Figueroa, María L. Delprato, Mohammad-Reza Hajirezaei, Martín L. Mayta, Anabella F. Lodeyro, Adriana R. Krapp, Néstor Carrillo

**Author notes:** Present address: Centre for Research in Agricultural Genomics (CRAG) Consorci CSIC-IRTA-UAB-UB Edifici CRAG Campus de Bellaterra de la UAB, Cerdanyola del Valles, Barcelona 08193, Spain. Present address: Centro para la Investigación en Ciencias de la Salud, Facultad de Ciencias de la Salud, Universidad Adventista del Plata, 25 de Mayo 99, E3103XAF, Libertador San Martín, Entre Ríos, Argentina. Correspondence to: Rocío C. Arce,; Adriana R. Krapp,; Néstor Carrillo.

## Abstract

Light significantly influences plant stress responses, with chloroplasts playing a pivotal role as both energy providers and light sensors. They communicate with the nucleus through retrograde signals, including secondary metabolites and reactive oxygen species (ROS). To investigate the contribution of chloroplast redox biochemistry to biotic responses, we studied the interactions of tobacco leaves expressing the alternative electron shuttle flavodoxin with virulent and nonhost *Pseudomonas syringae* pathovars under light and dark conditions. Flavodoxin is reported to limit light-dependent ROS propagation and over-reduction of the photosynthetic electron transport system under stress. Light intensified the hypersensitive response against the nonhost pathovar *tomato (Pto)*, but slowed disease progression caused by the virulent pathovar *tabaci (Pta)*. Flavodoxin mitigated light responses during both interactions, including decreased ROS levels, reduced stromule occurrence, and lower phytoalexin production, with different signatures depending on the pathovar. Similar leaf metabolic profiles were observed in the dark for both strains, with a general up-regulation of sugars, metabolic intermediates, and amino acids. In the light, instead, *Pta* increased sugars and intermediates, while *Pto* decreased them. Our results suggest that HR-like responses are elicited in the light even during virulent interactions, and that light effects are related to signals originating at the photosynthetic machinery.

**Highlights:**

- Light inhibits disease progression during a tobacco-*Pseudomonas* virulent interaction.
- Light exacerbates the hypersensitive response (HR) during a nonhost interaction.
- HR-like responses are elicited in the light even during virulent interactions.
- Plastid-targeted flavodoxin decreases plant damage only in photoperiod.
- The chloroplast redox state modulates plant biotic response.

## 1. Introduction

In the course of their life cycles, plants are continuously challenged by phytopathogenic bacteria, fungi, oomycetes and viruses. Upon pest attack, morphological and metabolic responses are activated to strengthen external barriers, set up defensive pathways and develop a toxic environment around the invading microorganism (Moreau et al., 2020). The final outcome of these biotic onslaughts, disease or resistance, depends on a number of factors, including the intrinsic diversity of plant-pathogen interactions, the type of plant immunity elicited by each of them, and the lifestyle of the attacking microorganism, either biotrophic, which requires living plant cells to multiplicate, or necrotrophic, which kills host cells and feeds on their remains (Fonseca and Mysore, 2019; Harris et al., 2020; Panstruga and Moscou, 2020; Yuan et al., 2021a,b). Interactions can thus be virulent, nonhost-resistant (NHR), defined as that exhibited by all cultivars of a plant species against all pathovars (pv) of a given microorganism (Harris et al., 2020; Panstruga and Moscou, 2020), or host-resistant, which depend on the presence and expression of plant resistance (*R*) alleles and is limited to one or few cultivars against a defined pv (Gill et al., 2015). Both NHR and host resistance may generate a hypersensitive reaction (HR) involving localized cell death (LCD) at the site of infection, which opposes a barrier of dead cells to the spread of biotrophic pathogens (Yuan et al., 2021a,b). Although the two types of resistant responses (NHR and host resistance) are initiated by distinct activation mechanisms, they involve basically the same suite of responsive genes (Tao et al., 2003; Bonfig et al., 2006; Harris et al., 2020; Panstruga and Moscou, 2020). As a consequence of the dynamic nature of plant-pathogen interactions, microorganisms that are closely related in terms of genetic background can display contrasting modes of interaction with a plant species, from full resistance to disease (Panstruga and Moscou, 2020).

In addition to these biological determinants, biotic interactions are also strongly influenced by environmental conditions. Light, in particular, has been shown to enhance plant responses during both NHR and host resistance, and to be required for full manifestation of the HR and LCD (Genoud et al., 2002; Zeier et al., 2004; Chandra-Shekara et al., 2006; Lukan et al., 2023). Even during virulent interactions such as that between tobacco and *Pseudomonas syringae* pv *tabaci* (*Pta*), the causative agent of tobacco wildfire disease, light provided partial protection against photosynthetic inactivation and suppressed bacterial growth in leaves (Wang et al., 2010; Cheng et al., 2016a,b). In many cases, the effect of illumination has been associated to increased generation of partially reduced and/or activated oxygen derivatives such as singlet oxygen (^1^O_2_), superoxide (O_2_^.-^) and hydrogen peroxide (H_2_O_2_), collectively known as reactive oxygen species (ROS). While the apoplast is regarded as the canonical site of the oxidative burst elicited by pathogen challenges (Bleau and Spoel, 2021), chloroplasts and peroxisomes produce the bulk of ROS in illuminated leaves under stress (Kuźniak and Kopczewski, 2020; Littlejohn et al., 2021), suggesting a possible contribution of the photosynthetic electron transport chain (PETC) to ROS propagation. Inactivation of photosynthesis is a universal feature of pathogen attacks (Cheng et al., 2016a,b), leading to over-reduction of the PETC and increased ROS synthesis, both of which may act in retrograde signaling to the nucleus (Dietz et al., 2019; Lodeyro et al., 2021).

The possible involvement of the PETC in the light responses to pathogens has been studied using photosynthetic inhibitors and redox-cycling agents (Cheng et al., 2016a). While the results obtained suggested a significant role of photosynthetic electron transport on light-dependent protection, the reagents employed had multiple impacts on plant physiology that might complicate interpretations. A complementary approach involves the use of alternative electron sinks that can affect the redox status of the chain and ROS production even in full illumination, and in a customized way. For instance, chloroplast targeting of a cyanobacterial flavodoxin (Fld), which accepts electrons at the reducing side of photosystem (PS) I and avoids PETC over-reduction, largely prevented ROS build-up and LCD during interactions of tobacco with the nonhost bacterium *Xanthomonas campestris* pv *vesicatoria* (Zurbriggen et al., 2009; Pierella Karlusich et al., 2017), and the necrotrophic fungus *Botrytis cinerea* (Rossi et al., 2017).

In this study, we examined the effects of Fld expression on the progression of tobacco wildfire disease elicited by *Pta*, employing the quasi-isogenic strain *P. syringae pv tomato* (*Pto*) as a nonhost counterpart. Lesions compatible with LCD were elicited by *Pto* inoculation in illuminated leaves, whereas those kept in the dark displayed only marginal symptoms. *Pta* inoculation, instead, led to leaf damage under both light and dark conditions, but significantly aggravated in the dark. Light- and dark-associated lesions were also qualitatively different: mostly chlorotic and necrotic, respectively. Pathogen-elicited ROS build-up and production of plastid stromules were higher under photoperiod, and particularly prominent during interaction with *Pto*. Accumulation of phytoalexins, instead, was stimulated under both light and dark conditions, especially by *Pto*. Determination of various soluble sugars, intermediate metabolites and amino acids revealed that metabolic responses were similar for the two pathovars in the dark, but differed significantly in the light. All these responses were mitigated by Fld presence only under photoperiod. The results suggest that light elicited HR-like lesions even during interaction with a virulent pathovar, thus providing some level of resistance, and that this effect was associated to the redox status of the PETC.

## 2. Materials and methods

### 2.1. Plant material

Preparation and characterization of homozygous tobacco plants (*Nicotiana tabacum* cv. Petit Havana) expressing a plastid-targeted Fld have been described previously (Tognetti et al., 2006). Briefly, the coding region of the *Anabaena* PCC7119 *fld* gene was fused in-frame to the 3’ end of a DNA sequence encoding the chloroplast transit peptide of pea ferredoxin-NADP^+^ reductase, and the fused construct was placed under the control of the constitutive cauliflower mosaic virus (CaMV) 35S promoter. Preparation and characterization of lines expressing a green fluorescent protein (GFP) targeted to chloroplasts have been reported by Köhler et al. (1997). The GFP coding sequence was fused in-frame to the 3’ end of a DNA fragment encoding the transit peptide of the Arabidopsis RecA protein, and the fused gene was cloned downstream of a double CaMV 35S promoter. Plants co-expressing Fld and GFP were generated by cross-pollination and selection of transformants to double homozygosity. Visualization of transgenic lines with strong and stable GFP fluorescence in chloroplasts was performed by confocal microscopy, whereas the presence and levels of Fld were determined by immunoblot assays (Tognetti et al., 2006; Ceccoli et al., 2012). Plants were grown at 250 µmol photons m^-2^ s^-1^ with a 16-h photoperiod, 28-20°C (day-night), and a relative humidity of 80%.

### 2.2. Inoculation with P. syringae pathovars

The *Pta* and *Pto* DC3000 strains were grown at 28°C in King’s B medium (KB; King et al., 1954) containing 50 μg mL^-1^ rifampicin (Rif). For the infiltration assays, fully expanded leaves (number five and six counting from the bottom) from 6-week-old tobacco plants were inoculated with a bacterial suspension containing 5 x 10^6^ colony forming units (CFU) mL^-1^ by using a needleless syringe. Leaves were also mock-infiltrated with 10 mM MgCl_2_ as control. For the dark condition, the inoculated leaves were covered with wooden paper envelopes.

### 2.3. Detection of stromules

Formation of stromules was evidenced in tobacco WT and *pfld* plants expressing chloroplast-targeted GFP, using a Zeiss 8000 laser scanning confocal microscope. Leaves inoculated with the various pathovars were infiltrated at 1 day post-inoculation (dpi) with 1 µM 4’,6-diamidino-2-phenylindole (DAPI), and incubated for an additional hour for nuclear visualization. Leaf discs were then collected, and fluorescence observed after excitation at 488 nm and emission between 505 and 530 nm. Stromules were quantified from maximum projections of z-stacks images using Fiji software (Schindelin et al., 2012). Stromule formation indexes were calculated as the ratio between the total number of stromules over the total number of chloroplasts in each field.

### 2.4. Detection of phytoalexins

Leaf discs were inoculated with ∼0.2 mL of mock or *P. syringae* suspensions, incubated for 16, 20, 26 and 42 h post-inoculation (hpi), and then exposed to UV light (∼302 nm). Images were taken with a Bio-Rad Gel Doc EQ Imaging System (Bio-Rad Laboratories, Hercules, USA). Fluorescence intensities were quantified with Fiji software taking a non-infiltrated region in each leaf analysed as blank (Schindelin et al., 2012).

### 2.5. RNA isolation and quantitative reverse-transcription (qRT)-PCR analysis

Total RNA was extracted from *P. syringae*- and mock-infiltrated leaf samples at 1 dpi, using the TriPure reagent (Sigma-Aldrich, St. Louis, USA), according to the manufacturer’s instructions, and reverse-transcribed with the MMLV enzyme (Invitrogen) and oligo(dT)_12-18_, using 1 µg of RNA as template. The qRT-PCR reactions were carried out in an AriaMx Real-time PCR System (Agilent), employing Platinum Taq DNA polymerase (Invitrogen) and SYBR Green I (Roche) to monitor the synthesis of double-stranded DNA. Relative transcript levels were determined using the delta threshold cycle (ΔΔC_t_) method (Schmittgen and Livak, 2008), in which each sample was normalized against the levels of tobacco elongation factor 1a (Schmidt and Delaney, 2010), and the levels of mock-inoculated samples. Primers (Supplementary Table S1) were designed using the “Primer3Plus” software (www.bioinformatics.nl/primer3plus/) with an annealing temperature of 55°C.

### 2.6. Analytical procedures

To determine electrolyte leakage, tobacco leaves were infiltrated with *P. syringae* strains. At 0, 1 and 2 dpi, 0.5-cm^2^ leaf discs were collected from the inoculated areas and incubated with distilled water for 30 min. Ion leakage was measured as the increase in conductance of the medium using a Horiba B-173 conductivity meter. At the end of the assay, samples were autoclaved to disrupt all cells, and total electrolyte contents were determined in the resulting solution.

To quantify the presence of hydroperoxides (-OOH), expressed as H_2_O_2_ equivalents, the FOX II assay (DeLong et al., 2002) was used in cleared leaf extracts from inoculated plants at 1 dpi, in parallel to mock controls. Leaf tissue corresponding to 4 cm^2^ was ground in liquid nitrogen and extracted with 300 µL of an 80:20 ethanol/water mixture containing 0.01% (w/v) butylated hydroxytoluene (BHT). After centrifugation at 3,000 *g* for 10 min, 15 µL of the supernatant were combined with 15 µL of 10 mM Tris–phenyl phosphine (TPP, a peroxide reducing agent) or the same volume of methanol, and incubated for 30 min at 25°C. Nine-hundred µL of FOX reagent (100 µM xylenol orange, 4 mM BHT, 250 µM ferrous ammonium sulphate, 25 mM H_2_SO_4_ in 90% methanol) were then added to each sample, and the absorbance at 560 nm was recorded 10 min after reagent addition. The absorbance differences between equivalent samples with or without TPP indicate the levels of peroxides, which were calculated using an H_2_O_2_ standard curve.

### 2.7. Measurements of soluble sugars, amino acids and metabolic intermediates

Metabolites were determined essentially as described by Ghaffari et al. (2016). Numerical analysis and quantification of individual compounds were carried out using the Empower Pro software (Waters, Milford, MA, USA) and authentic standards, respectively. Soluble sugars were determined by enzyme-coupled assays (Ghaffari et al., 2016). Metabolite separation and detection were carried out with an ion chromatography system (Dionex Thermofisher) connected to a triple quadrupole mass spectrometer QQQ6490 (Agilent Technologies), followed by ESI-MS/MS analysis (Ghaffari et al., 2016).

For amino acid determinations, samples were derivatized using the fluorescing reagent 6-aminoquinolyl-N-hydroxysuccinimidylcarbamate (AQC). Three mg of home-made AQC (IPK, Germany) were dissolved in 1 mL acetonitrile and incubated for 10 min at 55°C. The reagent was stored at 4°C and used for up to 1 week. For derivatization of the sample, 10 μL of the reagent solution were employed for each sample which contained 0.8 mL of 0.2 M boric acid, pH 8.8, and 10 μL of extract. Separation of soluble amino acids was performed by a newly developed method using ultra pressure reversed-phase chromatography (UPLC) AcQuity H-Class (Waters). The UPLC system consisted of a quaternary solvent manager, a sample manager-FTN, a column manager and a fluorescent detector (PDA eλ Detector). Separation was carried out on a C18 reversed-phase column (ACCQ Tag Ultra C18, 1.7 μm, 2.1 × 100 mm) with a flow rate of 0.7 mL per min and a duration of 10.2 min. The column was heated at 50°C during the whole run. Detection wavelengths were 266 nm for excitation and 473 nm for emission. The gradient was accomplished with four solutions prepared from two different buffers: eluent A concentrate and eluent B for amino acid analysis (Waters). Eluent A was pure concentrate, eluent B was a mixture of 90% LCMS water (Chemsolute) and 10% eluent B concentrate, eluent C was pure eluent B concentrate and eluent D was LCMS water. The column was equilibrated with a mixture of eluent A (10%) and eluent C (90%) for at least 30 min. The gradient was generated as follows: 0 min, 10% eluent A and 90% eluent C; 0.29 min, 9.9% A and 90.1% C; 5.49 min, 9% A, 80% B and 11% C; 7.1 min, 8% A, 15.6% B, 57.9% C and 18.5% D; 7.3 min, 8% A, 15.6% B, 57.9% C and 18.5% D; 7.69 min, 7.8% A, 70.9% C and 21.3% D; 7.99 min; 4% A, 36.3% C and 59.7% D; 8.68 min, 10% A, 90% C; and 10.2 min, 10% A and 90% C.

### 2.8. Statistical and graphical analyses

Data were analysed using one-way ANOVA and Duncan multiple range tests (*P* < 0.05). When the normality and/or equal variance assumptions were not met, Kruskall–Wallis one-way ANOVA between lines and Dunn’s multiple range tests were used (*P* < 0.1). Graphs were generated in R language (http://www.r-project.org/) using the ggplot2 library with the ‘geom_bar’ function for plotting bar graphs. The ‘heatmap.2’ function from the gplots library was used for plotting heat map graphs. For the latter, we standardized the dataset containing metabolite measures, to ensure that the data variables were on a comparable scale. Standardization was performed using the ‘scale’ function in R language, which centers and scales each variable, so that they had a mean of 0 and a standard deviation of 1.

## 3. Results

### 3.1. Illumination affects the nature and progress of tissue damage during the interaction of tobacco leaves with P. syringae pathovars

Young fully expanded leaves from WT tobacco plants were infiltrated with suspensions containing 5 x 10^6^ CFU mL^-1^ of *Pta* or *Pto*. Bacteria were inoculated at both sides of the central vein, one half of the leaf was covered to simulate dark conditions, and the plants were kept in the growth chamber with a 16/8 h photoperiod (see Material and Methods). Figure 1 shows typical results obtained at 4 dpi. Infiltration of illuminated leaves with *Pta* elicited symptoms consistent with wildfire disease (Figure 1a). Damage was significantly aggravated in the leaf regions kept in the dark, which displayed extended moist necrosis and evidence of tissue maceration, whereas dry chlorotic lesions were observed in illuminated leaves (Figure 1a). Exposure to *Pto* under normal photoperiod conditions led to lesions resembling LCD associated to the HR, and only marginal damage in the dark (Figure 1b), confirming the light dependence of this response (Genoud et al., 2002; Zeier et al., 2004; Chandra-Shekara et al., 2006; Lukan et al., 2023). A time-course study revealed that lesions were first detected at 3 dpi in leaf regions infiltrated with *Pta*, under both light and dark conditions (Supplementary Figure S1a), with more damage in the latter case. Light-dependent symptoms elicited by *Pto* were evident at 2 dpi (Supplementary Figure S2a).

**Figure 1:**
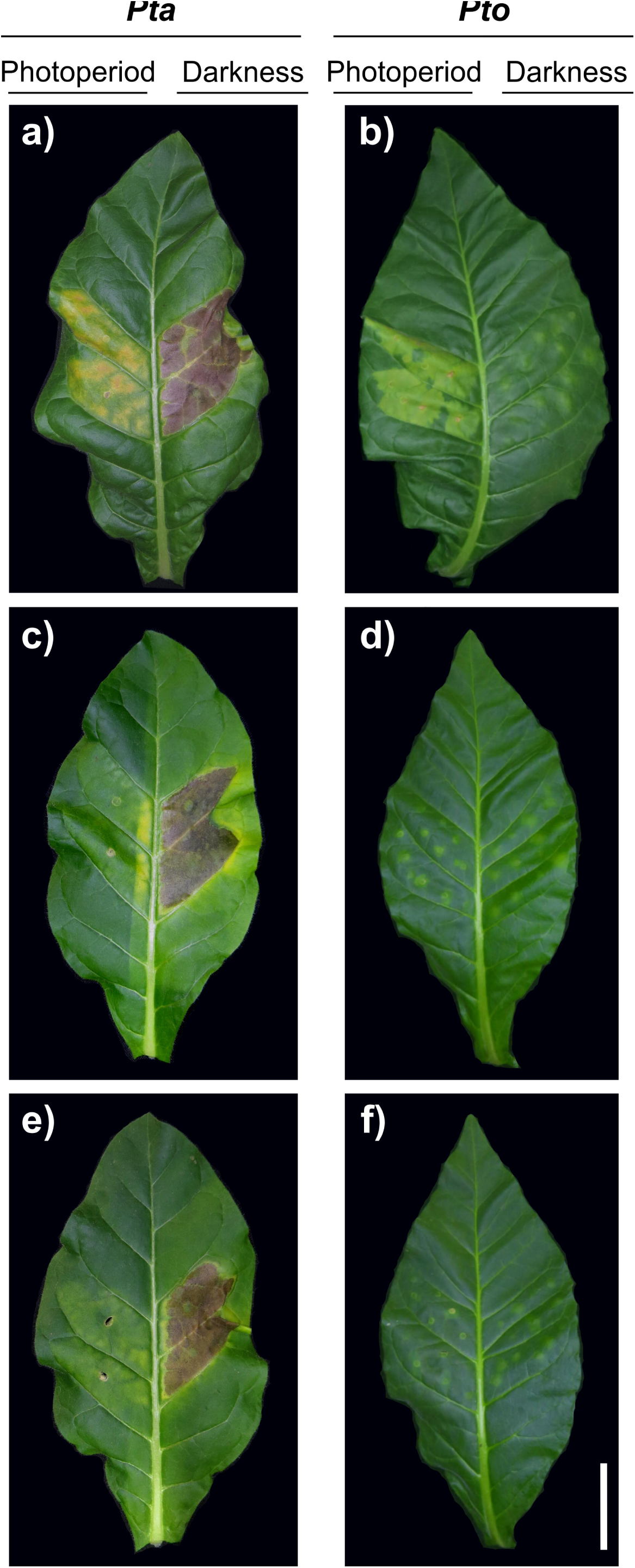
Effect of light during both virulent and resistant interactions between tobacco and *P. syringae* pathovars. Phenotypes of WT (a,b), *pfld*4-2 (c,d) and *pfld*5-8 (e,f) plants infected with *Pta* (a,c,e) or *Pto* (b,d,f) under photoperiod and dark conditions in the same leaf after 4 dpi. Bar = 10 cm.

To investigate if the light effect was associated to photosynthetic activity, the redox state of the PETC under illumination was manipulated by introduction of the alternative electron shuttle Fld (Lodeyro et al., 2021). As expected, the presence of the flavoprotein ameliorated lesions caused by *Pto* under photoperiod (Figure 1d,f; Supplementary Figure S2b,c), in good agreement with previous reports on other nonhost interactions (Zurbriggen et al., 2009). Surprisingly, Fld also limited the damage caused by *Pta* in illuminated leaf regions, even though the HR is not reported to occur in this virulent interaction (Figure 1c,e; Supplementary Figure S1b,c). Its introduction had no effect on the lesions caused by this strain in the dark (Figure 1c,e; Supplementary Figure S1b,c).

The degree of cell injury during *Pta* and *Pto* challenge was estimated by measuring the release of electrolytes at different times post-infiltration in both light and dark conditions. Determinations were extended to 3 dpi, when visual symptoms were apparent in WT plants for the two pathovars (Supplementary Figure S1). In line with the phenotypic results, Fld expression provided some protection under photoperiod for both pathovars, but not in the dark (Supplementary Figure S3).

### 3.2. Chloroplast Fld partially prevented light-dependent ROS accumulation in infiltrated leaves

The production of ROS, termed “oxidative burst”, is a landmark of successful pathogen recognition, and is required to fully activate plant defenses (Camejo et al., 2019; Kuźniak et al., 2020), including LCD (van Breusegem and Dat, 2006; Das and Roychoudhury, 2014). We therefore evaluated ROS production in leaves infiltrated with *Pta* or *Pto* under photoperiod and dark conditions. Since ROS build-up is one of the earliest cellular responses to pathogen challenge, samples were analysed at a pre-symptomatic stage (1 dpi) to visualize biochemical responses occurring before extensive tissue destruction. Leaves infiltrated with a saline solution (10 mM MgCl_2_) were used as a mock control. Quantification of total hydroperoxides (–OOH) in WT leaf extracts revealed a significant increase of ROS levels compared to mock-inoculated siblings in both interactions (Figure 2), and preferentially in the light (Figure 2). Fld presence prevented the ROS burst in illuminated leaves, but not in those kept in the dark (Figure 2).

**Figure 2:**
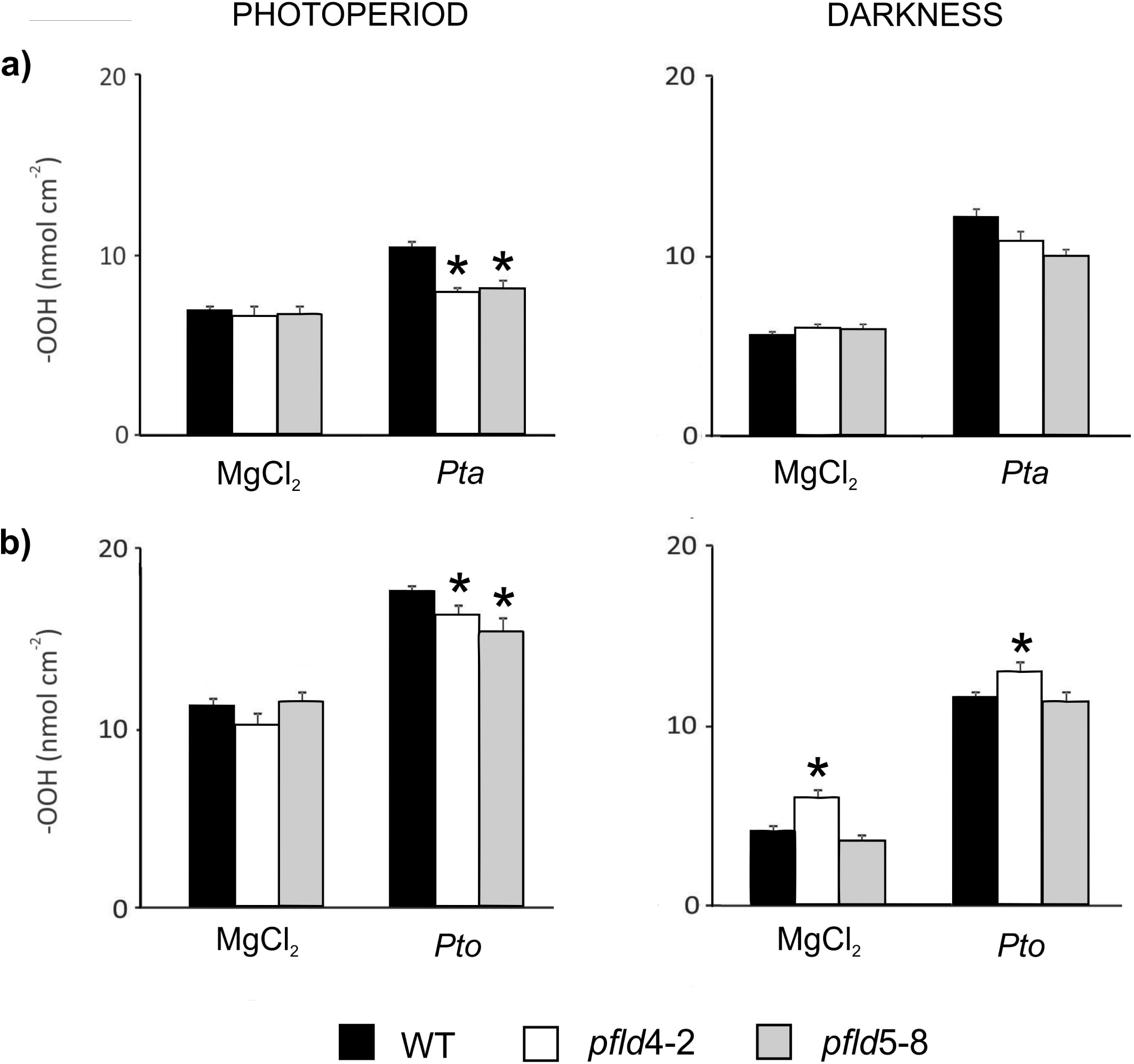
Hydroperoxide build-up during virulent and resistant interactions between tobacco and *P. syringae* pathovars. Total hydroperoxides (–OOH) were determined in WT and *pfld* lines infiltrated with *Pta* (a) or *Pto* (b) under photoperiod or dark conditions at 1 dpi. Data showed are means ± SE of 6 biological replicates per line corresponding to two independent experiments. Asterisks indicate statistically significant differences (*P <* 0.05) determined using one-way ANOVA and Duncan’s multiple comparison test.

### 3.3. Light increased stromule formation during both Pta and Pto interactions

Communication between chloroplasts and the nucleus in response to various environmental cues, including biotic stresses, may be mediated by stromules (stroma-filled tubular plastid extensions) which provide an additional conduit for transfer of a wider range of signalling molecules bypassing the cytosol (Mullineaux et al., 2020). Up-regulation of stromule formation has been associated with LCD during NHR and host resistance, and can facilitate LCD progression as part of the plant immune responses (Caplan et al., 2015). Their role during virulent interactions is much less known. We therefore evaluated stromule formation by confocal microscopic analysis of leaves expressing a chloroplast-targeted GFP, after inoculation with *Pta* or *Pto*. Images were taken at a pre-symptomatic stage (1 dpi). The results showed an increase of stromules in illuminated WT leaves during interactions with *Pta*, although markedly below the response elicited by the nonhost pathovar (Figure 3a). Estimations of the stromule formation index (SFI), calculated as the ratio of total number of stromules over total number of chloroplasts per image, provided quantitative support to this observation (Figure 3b). Expression of plastid-targeted Fld had little effect on *Pta*-induced stromule formation, while significantly inhibiting that elicited by *Pto* (Figure 3). In the dark, the formation of stromules was not significant in any of the lines or interactions assayed (Supplementary Figure S4).

**Figure 3:**
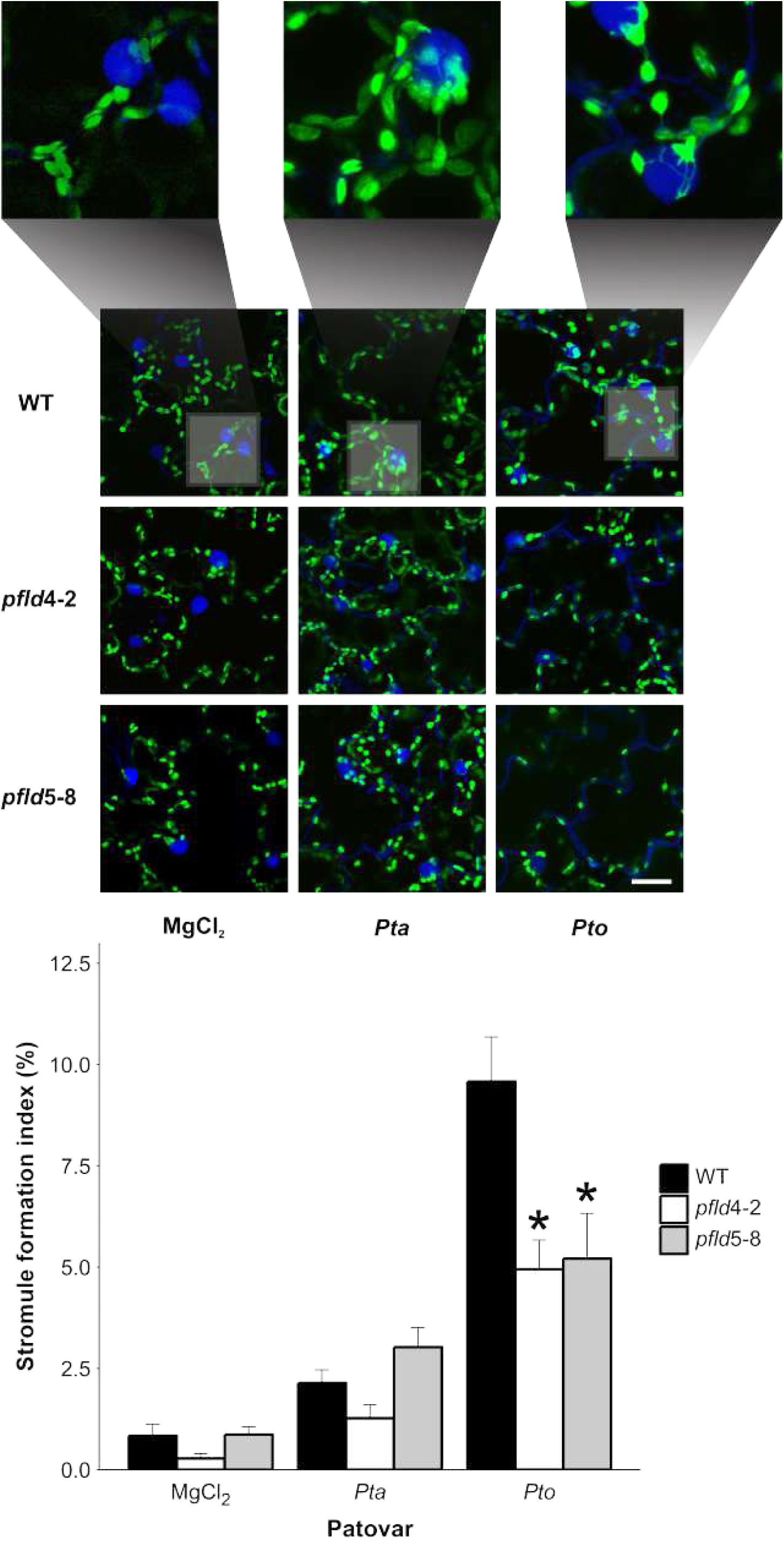
Induction of stromules during inoculation with *P. syringae* pathovars under photoperiod. (a) Typical micrographs showing stromule formation in epidermal tissue of WT and *pfld* lines infected with *Pta* or *Pto*. Tobacco plants expressing chloroplast-targeted GFP were kept under photoperiod, stained with 1 µM DAPI, and visualized at 1 dpi. Bar = 25 µm. (b) The stromule formation index (SFI) was determined as described in Materials and Methods. Data shown are means ± SE of 8-10 biological replicates per genotype corresponding to 2 independent experiments. Asterisks indicate statistically significant differences (*P* < 0.1) determined using Kruskall-Wallis test.

### 3.4. Biotic stress-dependent phytoalexin production is limited by chloroplast Fld under photoperiodic conditions

Secondary metabolites such as phytoalexins are part of the induced defense mechanisms against biotic stressors. Phytoalexins exhibit blue fluorescence under UV illumination, which can be used to monitor their accumulation during biotic interactions (Agati et al., 2022). Infiltrated WT leaves showed blue fluorescence around the infection zone under dark or light conditions for both *Pta* and *Pto*, whereas fluorescence was null in mock-inoculated leaves (Figure 4a). Phytoalexin accumulation proceeded at a slower rate in response to *Pta* compared to *Pto.* The blue fluorescence intensity was maximal for the latter microorganisms at 16 hpi and declined thereafter, whereas fluorescence of *Pta*-inoculated plants was undetectable until 26 hpi and was still increasing at 42 hpi (Figure 4b). Fluorescence intensities were slightly higher in the light for the two pathovars (Figure 4b).

**Figure 4:**
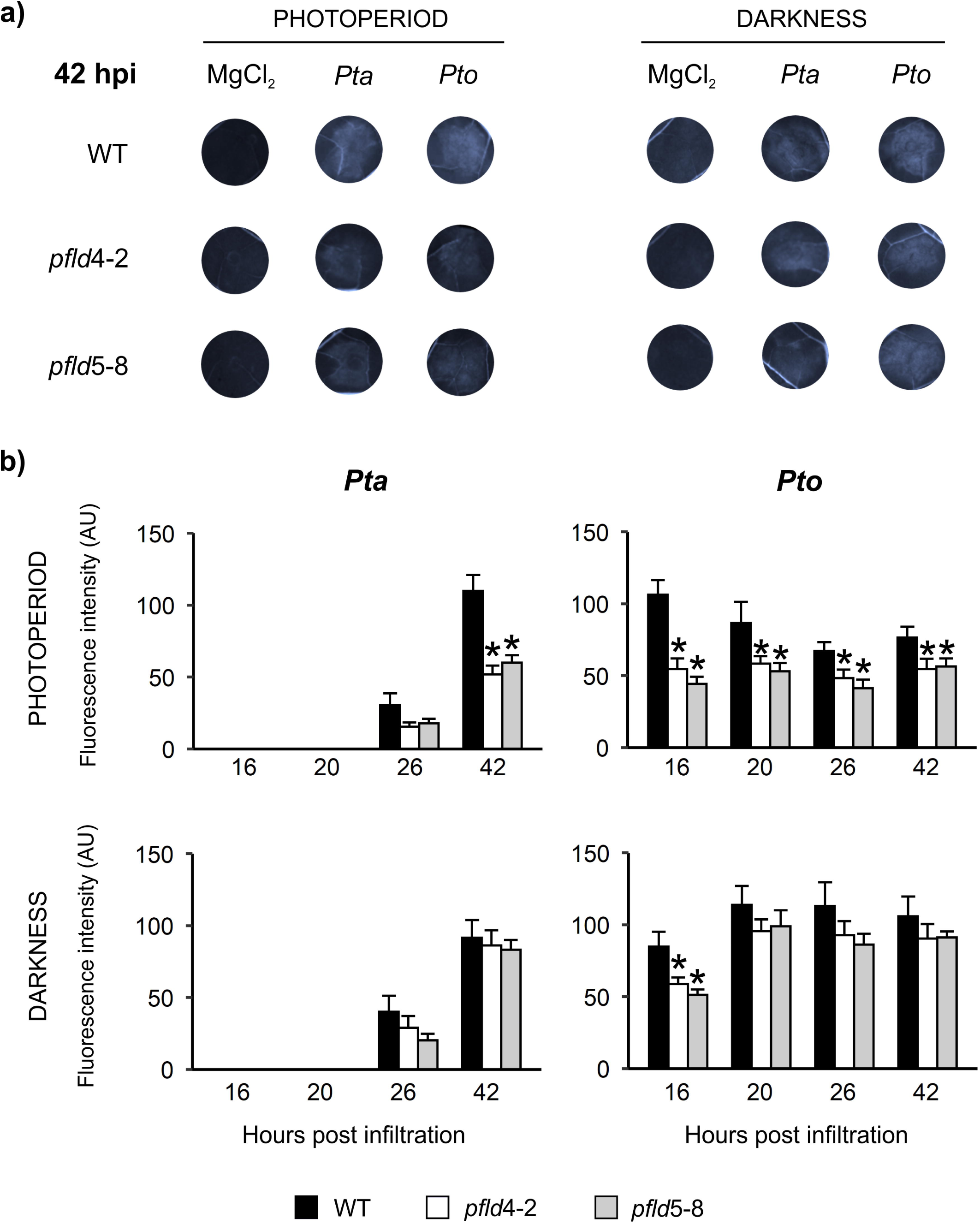
Phytoalexin accumulation in response to biotic interactions. The blue fluorescence was determined in WT and *pfld* leaves at various times after infection with *Pta* or *Pto*, under photoperiod and dark conditions, as described in Material and Methods. (a) Representative leaf discs infiltrated with *Pta* or *Pto* at 42 hpi under UV. (b) Quantification of fluorescence intensity at various hpi under photoperiod or darkness. Data shown are means ± SE of 3-4 biological replicates per genotype corresponding to 3 independent experiments. Asterisks indicate statistically significant differences (*P <* 0.05) determined using one-way ANOVA and Duncan’s multiple comparison test.

Phytoalexin production was lower in leaves from two independent *pfld* lines compared to their WT siblings when plants were incubated in the light, whereas no major differences were detected in the dark (Figure 4b).

### 3.5. Pathogenesis-related genes are induced in response to Pta infection

PR proteins have been widely implicated in plant resistance against a wide range of pathogens (Zribi et al., 2021). Their expression is generally induced during NHR and host-resistant interactions and less in virulent processes (Kaur et al. 2020). To determine if this induction was affected by light or the genotype, we determined the transcript levels of three PR genes in *Pseudomonas*- and mock-infiltrated plants: (*i*) *9LOX*, which encodes a 9S-lipoxygenase involved in lipid peroxidation during plant responses to pathogen infection (Rustérucci et al., 1999); (*ii*) *PR1b*, known to be part of core components of plant defense (Iqbal et al., 2021); and (*iii*) the ethylene-responsive transcription factor *Pti5*, which contributes to basal resistance and is induced during NHR (Pierella Karlusich et al., 2017). These genes were chosen because they belong to different PR families and participate in different aspects of defense (Zribi et al., 2021).

Expression of the three marker genes was induced by *Pto* infiltration under both light and dark regimes (Supplementary Figure S5). Fld presence had little effect on this up-regulation, except for *9LOX*, whose induction was lower for the transformants under both conditions compared to WT siblings, and *PR1b*, which displayed lower induction under dark in the *pfld* lines (Supplementary Figure S5). Despite the virulent nature of the interaction, marker genes were also induced after *Pta* challenge, with fold changes ranging between 4 and 1000 (Supplementary Figure S5). Light effects were moderate, except for *PR1b*, which was ∼3-fold less induced in the light than in the dark (Supplementary Figure S5). Fld introduction decreased the induction of *9LOX* under both regimes, whereas it caused a moderate up-regulation of *PR1b* and *Pti5*, especially in photoperiod (Supplementary Figure S5).

### 3.6. Central metabolic routes were differentially affected by the biotic stress depending on the illumination conditions

The multiple mechanisms involved in the plant responses to biotic stresses imply a reprogramming of central metabolic routes and a massive redistribution of resources, from growth to defense (Iqbal et al., 2021). To determine which type of metabolic adjustments take place in WT and *pfld* leaves infiltrated with *Pta* or *Pto*, we measured the levels of soluble carbohydrates, intermediate metabolites and amino acids at 1 dpi under photoperiod and dark conditions.

Sugars play significant roles throughout the plant life cycle as nutrients and also as signal molecules. Levels of glucose, fructose and sucrose were already ∼2-fold higher in the light than in the dark for mock-treated WT leaves (Figure 5), likely reflecting increased production of photosynthates under illumination. Treatment with *Pta* led to further increases of the three sugars in the light (Figure 5a). Glucose and fructose contents were also up-regulated by *Pta* in the dark, whereas sucrose levels were not modified (Figure 5b). Mock-inoculated plants expressing Fld contained less free glucose and fructose than their WT counterparts under both light and dark, and this trend was maintained even after *Pta* treatment (Figure 5a,b). Sucrose levels were instead not modified by the infection in *pfld* leaves incubated in the light, and decreased slightly in the dark (Figure 5a,b).

**Figure 5:**
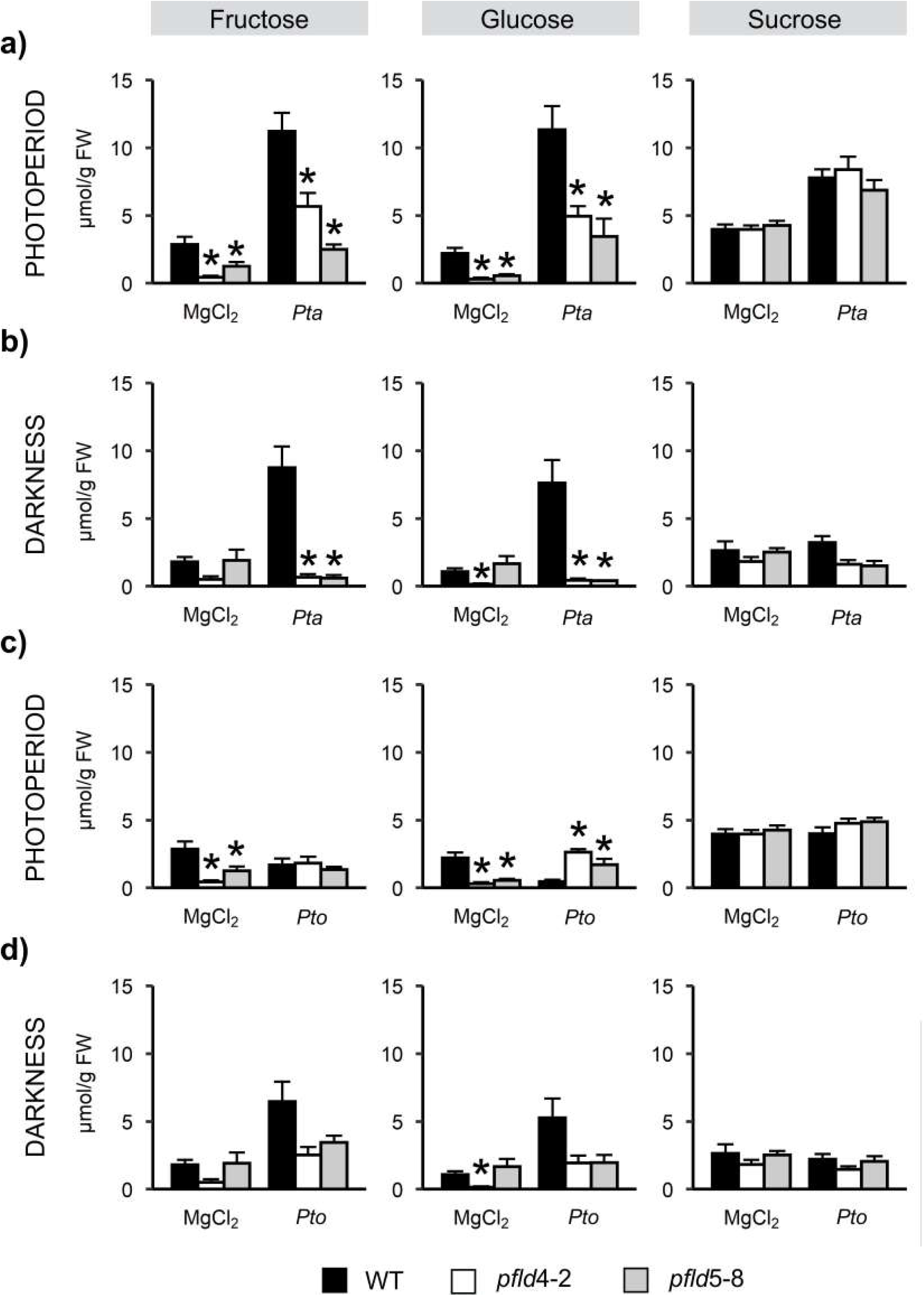
Sugar contents in plants inoculated with *Pta* or *Pto* under light or dark regimes. Levels of carbohydrates in photoperiod (a,c) and darkness (b,d) for the virulent *Pta* interaction (a,b) and for the nonhost *Pto* pathovar (c,d). Data shown are means ± SE of 6 biological replicates per genotype. Asterisks indicate statistically significant differences (*P <* 0.05) determined using one-way ANOVA and Duncan’s multiple comparison test.

Exposure of illuminated WT plants to *Pto* failed to increase glucose and fructose as in *Pta*-treated leaves, leading instead to moderate declines in the contents of the two carbohydrates (Figure 5c). These decreases were fully prevented (fructose), or even offset (glucose) by Fld expression (Figure 5c). Sucrose contents were not affected by either Fld or pathogen challenge (Figure 5c). In the dark, plant responses to *Pto* resembled those elicited during infection with *Pta* under the same conditions (Figure 5d).

The leaf contents of 24 intermediate metabolites were determined for the various conditions, pathovars and genotypes. Results are summarized as heatmaps in Figure 6, following normalization (see Materials and methods) with detailed data provided in Supplementary Figures S6-S9. A key trend observed in the heatmaps was that, under light conditions, the interaction with the virulent pathovar *Pta* led to an overall increase in metabolic intermediates relative to mock-infiltrated plants (Figure 6a). In contrast, the interaction with the nonhost pathovar *Pto* resulted in a decrease of most metabolites compared to the mock treatment (Figure 6b). Under dark conditions, both pathovars resulted in a general increase in metabolic intermediates across WT and transgenic tobacco lines, suggesting a more uniform metabolic response in the absence of light.

**Figure 6:**
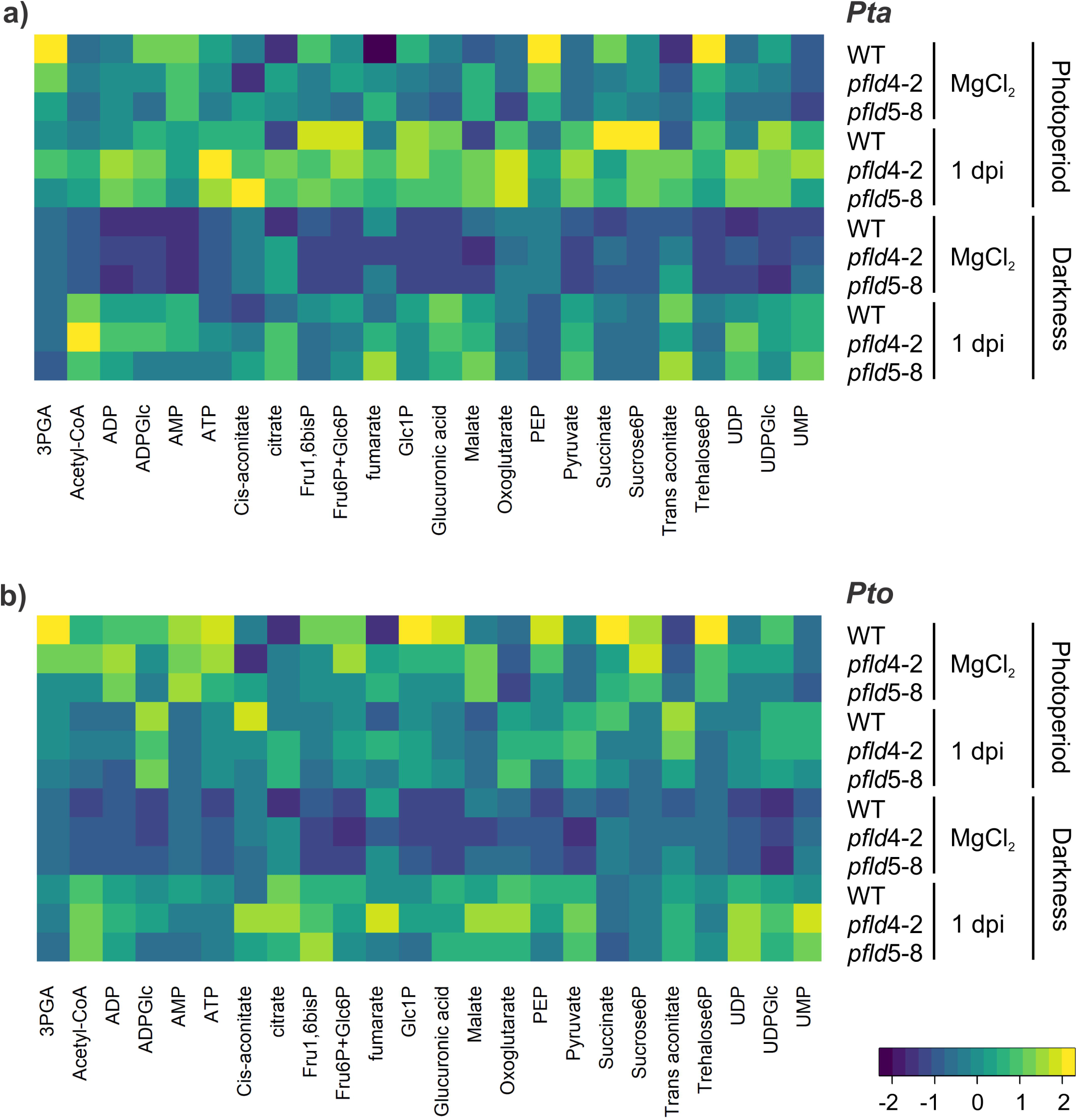
Relative contents of intermediate metabolites during biotic interactions under light and dark conditions. Heat maps of relative amounts of intermediate metabolites for treatment with *Pta* (a) or *Pto* (b). Each value for a compound was normalized based on the highest and lowest values observed across all lines and conditions for each pathovar. The colour scale corresponds to the standardized scores, green for low values and red for high values. Data shown are means of 6 biological replicates per genotype.

Analysis of individual metabolites indicates that most of them, including adenylates, accumulated to higher levels in illuminated mock-treated WT leaves compared to dark-adapted counterparts (Supplementary Figures S6-S9). Infiltration with *Pta* led to moderate changes in the light, with increases in phosphorylated sugars, pyruvate, oxoglutarate, succinate, fumarate, glucuronic acid, UMP and UDP-glucose, and down-regulation of 3-phospho-glyceric acid (3-PGA), phospho-*enol*-pyruvate (PEP) and trehalose 6-phosphate (Supplementary Figure S6). When the same treatment was carried out in the dark, a massive reprogramming occurred, with 16 out 24 assayed metabolites increasing upon infection and only one (PEP), decreasing (Supplementary Figure S7).

Fld expression had little impact in mock-inoculated leaves, but led to higher levels of some metabolites (adenylates, acetyl-CoA, UDP, intermediates of the tricarboxylic acid [TCA] cycle) relative to nontransformed siblings in *Pta*-infected leaves kept in the light (Supplementary Figure S6). Fld presence had instead negligible effects in the dark, except for ATP and citrate, whose levels were up-regulated in the transformants, compared to WT controls (Supplementary Figure S7). Some of the metabolites that showed increased levels in illuminated *pfld* leaves (ADP, UDP, acetyl-CoA) are involved in multiple metabolic pathways, and kept this trend after *Pta* challenge (Supplementary Figures S6, S7). Citrate and malate also showed higher contents in *pfld* leaves, which might indicate increased flux through the TCA cycle in these plants. Succinate was instead decreased (Supplementary Figure S6), suggesting a potential pathway that is consuming this compound in Fld-expressing lines.

A comparison with the response of WT plants against *Pto* revealed significant differences. Fourteen out of 24 metabolites were down-regulated in the light, with only four (citrate, oxoglutarate and the aconitate isomers) increasing their levels upon infiltration (Supplementary Figure S8). The dark response was entirely different, with 18 increases and no decreases (Supplementary Figure S9; Figure 6b), strikingly resembling the one elicited against *Pta* (Supplementary Figure S7). Fld had little effect on these metabolic profiles, except for *cis*-aconitate decrease under illumination, and up-regulation in the dark (Supplementary Figures S8, S9). In summary, results of Supplementary Figures S6-S9, as well as the normalized heatmaps of Figure 6 show that metabolite profiles differed between lines in their response to light-dark treatments, and as a consequence of Fld expression, but the overall trends were similar for mock-infiltrated samples. Exposure to the microorganisms in the dark also led to equivalent global responses for both *Pta* and *Pto*, while they exhibited major differences in the light (Figure 6).

The great demand for carbon and energy sources caused by the activation of defenses during pathogen challenge usually affects amino acid metabolism, driving them towards energy-generating pathways. We thus measured the levels of 17 amino acids in WT and *pfld* plants infiltrated with both pathovars, under photoperiod or in darkness. Figure 7 shows that there was a differential response to *Pta* and *Pto* in the light, similar to that observed for intermediate metabolites. Inoculation with *Pta* in the light resulted in increased levels of 10 out of 17 assayed amino acids (Supplementary Figure S10). Dark incubation rendered similar results (levels of 15 amino acids were up-regulated), with nine in common between the two regimes, notably Tyr (Supplementary Figure S11; Figure 7a). Ala, Gly and His were the only amino acids whose levels decreased upon *Pta* treatment in the light, whereas they increased in the dark (Supplementary Figures S10, S11). Fld presence had little effect on the amino acid levels during *Pta* infection in the light, except for the increase of Pro and Gly (Supplementary Figure S10), but led to significant increases in 15 of them in the dark (Supplementary Figure S11; Figure 7a). It is worth noting, within this context, that some of these amino acids (Ala, Arg, Ile, Leu, Pro) were already higher in mock-infiltrated *pfld* plants compared to WT counterparts, and this trend was maintained after *Pta* treatment (Supplementary Figure S11).

**Figure 7:**
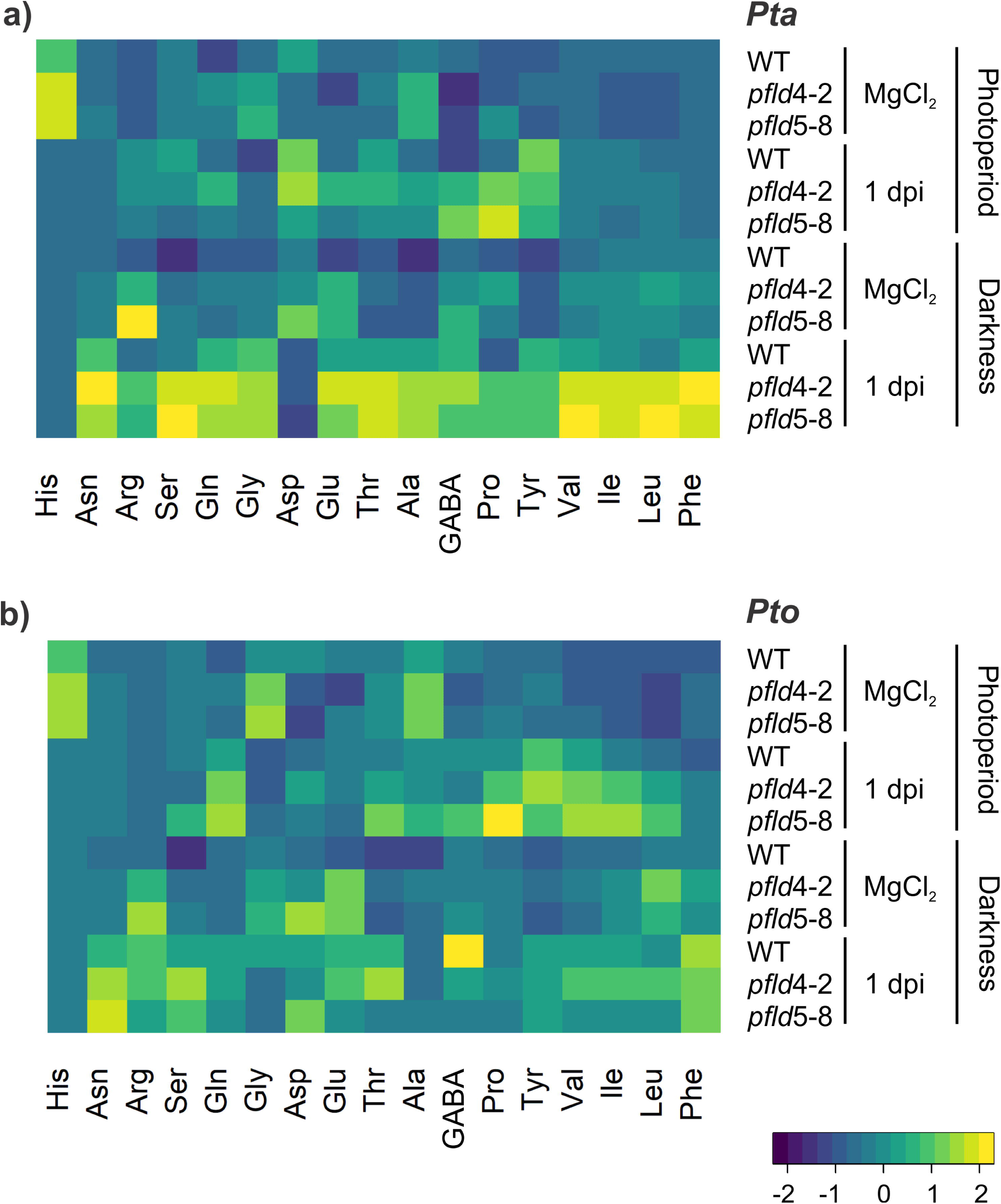
Relative contents of amino acids during biotic interactions under light and dark conditions. Heat maps of relative amounts of amino acids for treatments with *Pta* (a) or *Pto* (b). Each value for a compound was normalized based on the highest and lowest values observed across all lines and conditions for each pathovar. The colour scale corresponds to the standardized scores, green for low values and red for high values. Data shown are means of 6 biological replicates per genotype.

The light response of WT plants to *Pto* was moderate, with 8 amino acids increasing their levels (Asn, GABA, Gln, Ile, Leu, Pro, Tyr, Val), and two (Gly and His) declining (Supplementary Figure S12), whereas the dark response involved up-regulation of virtually all assayed amino acids (Supplementary Figure S13; Figure 7b). Fld enhanced further the accumulation of Gln, Leu, Ile, Val and Pro in *Pto*-infiltrated leaves in the light (Supplementary Figure S12), but had little effect in the dark, except for Pro (Supplementary Figure S13).

Therefore, analysis of individual amino acid profiles (Supplementary Figures S10-S13), and the normalized heatmaps of Figure 7 confirm the differential response to *Pta* and *Pto* in the light, similar to that observed for intermediate metabolites.

## 4. Discussion

Although plants have evolved an innate immune system to detect and cope with diverse biotic attacks, some species remain vulnerable to individual pathogenic microorganisms. Understanding the responses of these susceptible plants to pathogens is an area that requires further investigation (Gorshkov and Tsers, 2022). On the other hand, while the immune system provides a fundamental defense mechanism, the effectiveness of these protective barriers can be influenced by various factors, including environmental conditions. One crucial factor that significantly impacts plant defence is light, which not only provides an energy source via photosynthesis, but also signalling cues for developmental and environmental responses that influence multiple aspects of plant physiology (Iqbal et al., 2021). Although the role of illumination during plant-pathogen interactions is well documented (Delprato et al., 2015; Lukan et al., 2023), its relation with the photosynthetic machinery remains largely unexplored. We addressed this issue by altering the redox state of the PETC through the introduction of a cyanobacterial Fld targeted to plastids (Tognetti et al., 2006), and studying the response of these Fld-expressing plants during virulent and nonhost interactions with *P. syringae* pathovars.

The results showed that light had contrasting phenotypic effects on the two types of interactions. It enhanced the chlorotic lesions elicited by nonhost *Pto*, an effect that has been associated with full manifestation of the HR (Genoud et al., 2002; Zeier et al., 2004; Chandra-Shekara et al., 2006; Lukan et al., 2023), but ameliorated the damage caused by virulent *Pta* (Figure 1). Cheng et al. (2016a,b) reported that illumination led to protracted appearance of *Pta*-induced lesions. We confirmed these observations, although qualitative phenotypic differences were also evident. Incubation of the inoculated leaves in the dark led to extensive tissue maceration and brown necrosis, aggravating the typical symptoms of wildfire disease (Figure 1). In the light, instead, the *Pta*-induced lesions were mostly chlorotic, with few or no necrotic spots, a phenotype somehow intermediate between those elicited by *Pto* in the light and *Pta* in darkness (Figure 1).

While light can act at various levels and use different photosensors, the protective effect of Fld suggests an involvement of the PETC in these biotic responses. Fld introduction mitigated the light-dependent visual symptoms and cell damage in both interactions (Figure 1; Supplementary Figure S1-S3), but was inconsequential in the dark. The signalling role played by photosynthesis during pathogen challenges has been inferred from a number of observations (Zeier et al., 2004; Chandra-Shekara et al., 2006; Kuźniak and Kopczewski, 2020; Littlejohn et al., 2021; Yang and Luo, 2021; Yang et al., 2021), and could be mediated by chloroplast ROS and/or the redox state of PETC components. The photosynthetic machinery is an early and primary target of pathogens, resulting in photoinhibition, over-reduction of the PETC and delivery of the excess of energy and reducing power to oxygen (Kuźniak and Kopczewski, 2020; Littlejohn et al., 2021; Sun et al., 2021; Yang et al., 2021). Chloroplast ROS, which accumulate due to photosynthetic inactivation, might act as signalling molecules modulating plant responses during the biotic assault. The protective effect of Fld as an alternative electron shuttle of the PETC would concur with this possibility, as it prevents ROS build-up (Figure 2), although Fld can also modulate the redox state of chain intermediates that might participate in retrograde signaling by preventing their over-reduction (Gómez et al., 2020). Distinction between these possibilities requires further investigation.

Extensive research has focused on the formation of stromules, which have been found to be closely associated with the innate immune response. Stromules, induced by oxidative stress, establish direct connections between chloroplasts and the nucleus, bypassing the cytosol and facilitating communication between compartments and retrograde signalling (Caplan et al., 2015; Hanson and Hines, 2018; Kumar et al., 2018; Mullineaux et al., 2020). We observed that stromule formation was light-dependent for both microorganisms, with a 10-fold increase in SFI for *Pto* but just 2-fold for *Pta* (Figure 3; Supplementary Figure S4), indicating a weaker response in the virulent interaction (Figure 3b). Fld decreased stromule production, suggesting that *pfld* plants experienced lower stress pressure than WT siblings at this pre-symptomatic stage. Another critical aspect of host immunity is the production of phytoalexins, secondary metabolites known for their antimicrobial and antioxidant properties (Costet et al., 2002; Stringlis et al., 2019). The protracted build-up of phytoalexin levels in *Pta*-infected leaves, relative to *Pto*-inoculated siblings (Figure 4) agrees well with previous observations (Agati et al., 2022), confirming the weaker and delayed response of tobacco plants toward *Pta*, which ultimately leads to disease. Fld expression decreased the blue fluorescence intensity due to phytoalexins in the light, again highlighting the role of photosynthesis in defense. A similar effect has been reported in the context of necrotrophic fungal infection, where *pfld* plants exhibited lower phytoalexin accumulation compared to their WT counterparts (Rossi et al., 2017).

During biotic stresses, plants undergo significant metabolic reprogramming to effectively respond and defend against pathogenic attacks. This reprogramming involves complex changes in various pathways, allowing plants to allocate resources towards defense responses and strengthen their immune system (Iqbal et al., 2021). We evaluated the levels of sugars, amino acids and intermediate metabolites at 1 dpi and found different response patterns depending on the pathovars and illumination conditions (Figures 5-7; Supplementary Figures S6-S13). Glucose and fructose contents were significantly higher in leaves inoculated with the two pathovars and kept in the dark, with Fld presence partially preventing this increase (Figure 5). In the light, instead, *Pta* and *Pto* displayed sharply contrasting behaviours, with *Pta* eliciting hexose build-up as in the dark (Figure 5a,b), and *Pto* decreasing their levels (Figure 6c,d), presumably due to full manifestation of the HR under illumination (Genoud et al., 2002; Lukan et al., 2023), as carbohydrates would be required for defence deployment during this nonhost interaction.

There was also a clear accumulation of many intermediates of the glycolytic pathway in WT leaves exposed to *Pta* infection under both photoperiod and dark conditions (Supplementary Figures S6, S7; Figure 6a). ATP and ADP contents were also raised during the biotic stress in illuminated *pfld* plants. Metabolites related to the TCA cycle showed a similar behaviour to those of glycolysis in inoculated leaves, with a major accumulation in the presence of light (Supplementary Figure S6), except for acetyl-CoA, which was more prominent in the dark (Supplementary Figure S7). Virulent *Pta* increased the levels of several amino acids under light and dark, with a slightly higher trend in *pfld* lines (Supplementary Figures S10, S11; Figure 7a).

The metabolic reprogramming in response to *Pto* infiltration displayed different patterns compared to the virulent interaction. Although most glycolytic intermediates and amino acids accumulated in the dark after inoculation, some of them (3PGA, PEP, Gly, Ala) were higher in control conditions under photoperiod. Similar results were obtained for ATP and ADP in the light, with their levels decreasing upon *Pto* infiltration (Supplementary Figures S10-S13; Figure 7b). Intermediates of the TCA cycle, as well as amino acids potentially produced by this pathway, increased their relative levels in the dark, with the exception of succinate, Gln and Pro. In conclusion, metabolic responses elicited by the two pathovars were similar in the dark, but differed significantly in the light, with several sugars and metabolic intermediates increasing upon *Pta* infection but decreasing after *Pto* challenge. This might indicate that metabolic intermediates and precursors are actively redirected to building up defensive barriers in the latter interaction. Comprehensive models summarizing these observations are shown in Figure 8. Fld introduction had moderate effects, reflected by the differential increase of some compounds under the adverse situation (Figures 6-8; Supplementary Figures S6-S13).

**Figure 8:**
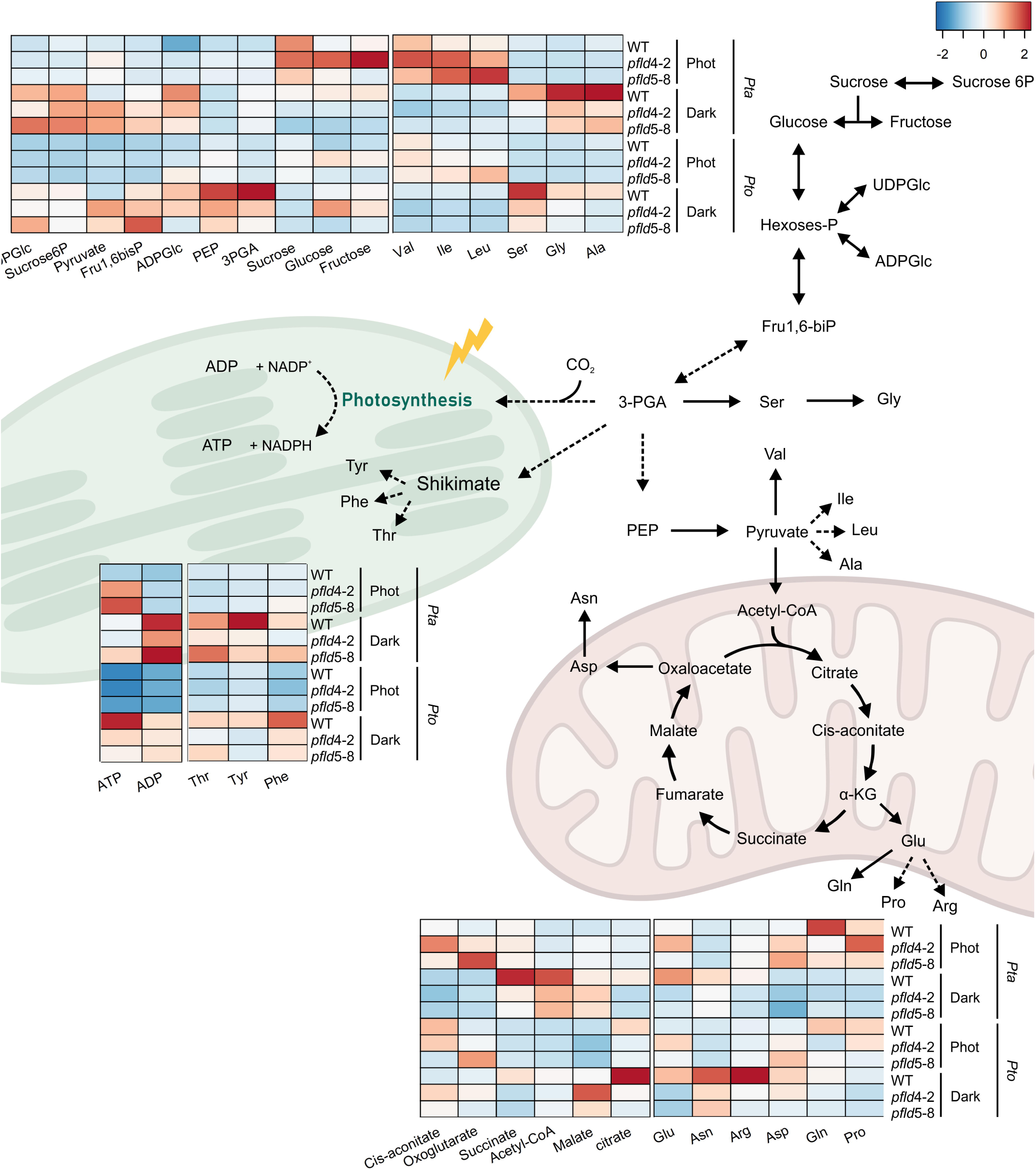
Metabolic reprograming in response to illumination conditions under *Pta* and *Pto* infection. Summary of metabolic networks behaviour of leaves facing the virulent (*Pta*) and NHR infection (*Pto*) under dark and light conditions for all tobacco lines. Quantitative data of this figure are described in Figures 5-7 and Supplementary Figures S6-S13. Metabolite levels are presented as the ratio of each measurement relative to the corresponding mock-inoculated controls (*Pta*/MgCl_2_ and *Pto*/MgCl_2_) under each condition. The resulting values were standardized using the ‘scale’ function in R, which centers and scales each variable to have a mean of 0 and a standard deviation of 1, ensuring comparability across metabolites. Positive values indicate increased levels under infiltration, while negative values indicate decreased levels. The color scale represents standardized scores, with blue indicating low values and red indicating high values. Data represent the means of six biological replicates per genotype.

Based on these observations, we propose that light triggers similar HR-type pathways in susceptible and resistant plants, although with different strength and efficiency, and that this is the source of the partial protection conferred by Fld to *Pta*-associated damage in the light (Fig. 1, Supplementary Fig. S1; see also Cheng et al., 2016a,b). We also propose that the PETC modulates this light response. Results obtained in this study provide insights into the role of light and the chloroplast redox status in the context of plant-pathogen interactions, and highlight the importance of light as a modulator of defense responses. They also illustrate the complexity of the relationship between light, chloroplast redox biochemistry and disease progression, suggesting that factors beyond ROS may contribute to the development of plant symptoms. Additionally, the study sheds light on the involvement of stromules and phytoalexins in the plant innate immune response and their modulation by pathogen infection and Fld expression. Moreover, the differential metabolic responses observed between the two pathogens indicate distinct metabolic reprogramming during biotic stress. Further investigations are needed to unravel the molecular mechanisms underlying these processes and to explore their usefulness in developing strategies for disease management and crop improvement.

## Author contributions

RCA, ARK and NC defined the goals of the research and designed pertinent experiments. RCA, MD, NF, MLD, MLM performed the experiments. MRH, AFL, ARK and NC obtained the funding. All authors participated in analysis, writing and editing the manuscript.

## Acknowledgements

The authors wish to thank Diego Aguirre for his technical assistance with plant growth at IBR. We also acknowledge to Rodrigo Vena for his technical assistant with the confocal microscopy at IBR.

## Legends to Supplementary material

**Supplementary Figure S1: Time course of tissue damage after infiltration of tobacco leaves with *Pta* under photoperiod and dark conditions.** Representative leaves from WT (a), *pfld*4-2 (b) and *pfld*5-8 (c) plants at 1, 2, 3 and 4 days post-infiltration (dpi) with *Pta*. Dashed lines indicate the inoculated areas. Bar = 10 cm.

**Supplementary Figure S2: Time course of tissue damage after infiltration of tobacco leaves with *Pto* under photoperiod and dark conditions.** Representative leaves from WT (a), *pfld*4-2 (b) and *pfld*5-8 (c) plants at 1, 2, 3 and 4 days post-infiltration (dpi) with *Pto*. Dashed lines indicate the inoculated areas. Bar = 10 cm.

**Supplementary Figure S3: Cell injuries caused by *P. syringae* pathovars.** Ion leakage of WT and *pfld* plants through time post-infection with *Pta* (a) or *Pto* (b) under photoperiod and dark conditions. Data showed are means ± SE of 6 biological replicates per genotype corresponding to two independent experiments. Asterisks indicate statistically significant differences (P < 0.05) determined using one-way ANOVA and Duncan’s multiple comparison test.

**Supplementary Figure S4: Stromule formation in response to *Pta* or *Pto* inoculation in the dark.** Representative images of WT and *pfld* lines expressing chloroplast-targeted GFP. Stromule formation in epidermal leaves of WT and *pfld* lines infected with *Pta* or *Pto*. Images are representative of 2 independent experiments. Bar = 25 µm.

**Supplementary Figure S5: Expression patterns of pathogenesis-related genes in tobacco leaves inoculated with *Pta* or *Pto* under photoperiod or dark conditions.** Data reported are means ± SE of 4 biological replicates per genotype. Asterisks indicate statistically significant differences (*P <* 0.05) determined using one-way ANOVA and Duncan’s multiple comparison test. 9LOX, 9S-lipoxygenase 5; PR1b, pathogenesis related protein 1b; Pti5, ethylene response factor Pti 5.

**Supplementary Figure S6: Contents of intermediate metabolites in WT and *pfld* plants exposed to *Pta* infection under photoperiod condition.** Data shown are means ± SE of 6 biological replicates per genotype. Asterisks indicate statistically significant differences (*P <* 0.05) determined using one-way ANOVA and Duncan’s multiple comparison test.

**Supplementary Figure S7: Contents of intermediate metabolites in WT and *pfld* plants exposed to *Pta* infection in the dark.** Data shown are means ± SE of 6 biological replicates per genotype. Asterisks indicate statistically significant differences (*P <* 0.05) determined using one-way ANOVA and Duncan’s multiple comparison test.

**Supplementary Figure S8: Contents of intermediate metabolites in WT and *pfld* plants exposed to *Pto* inoculation in the presence of light.** Data shown are means ± SE of 6 biological replicates per genotype. Asterisks indicate statistically significant differences (*P <* 0.05) determined using one-way ANOVA and Duncan’s multiple comparison test.

**Supplementary Figure S9: Contents of intermediate metabolite in WT and *pfld* plants exposed to *Pto* inoculation in the dark.** Data shown are means ± SE of 6 biological replicates per genotype. Asterisks indicate statistically significant differences (*P <* 0.05) determined using one-way ANOVA and Duncan’s multiple comparison test.

**Supplementary Figure S10: Amino acid levels in WT and Fld-expressing leaves exposed to *Pta* infection in the presence of light.** Data shown are means ± SE of 6 biological replicates per genotype. Asterisks indicate statistically significant differences (*P <* 0.05) determined using one-way ANOVA and Duncan’s multiple comparison test.

**Supplementary Figure S11: Amino acid levels in WT and Fld-expressing leaves exposed to *Pta* infection in the dark.** Data shown are means ± SE of 6 biological replicates per genotype. Asterisks indicate statistically significant differences (*P <* 0.05) determined using one-way ANOVA and Duncan’s multiple comparison test.

**Supplementary Figure S12: Amino acid levels in WT and Fld-expressing leaves exposed to *Pto* inoculation in the presence of light.** Data shown are means ± SE of 6 biological replicates per genotype. Asterisks indicate statistically significant differences (*P <* 0.05) determined using one-way ANOVA and Duncan’s multiple comparison test.

**Supplementary Figure S13: Amino acid levels in WT and Fld-expressing leaves exposed to *Pto* inoculation in the dark.** Data shown are means ± SE of 6 biological replicates per genotype. Asterisks indicate statistically significant differences (*P <* 0.05) determined using one-way ANOVA and Duncan’s multiple comparison test.

**Supplementary Table S1:** Primers used for qRT-PCR determinations.

