## Supplementary figures and images for "Light and chloroplast redox state modulate the progression of tobacco leaf infection by *Pseudomonas syringae* pv *tabaci*"

### Supplementary Figure S1

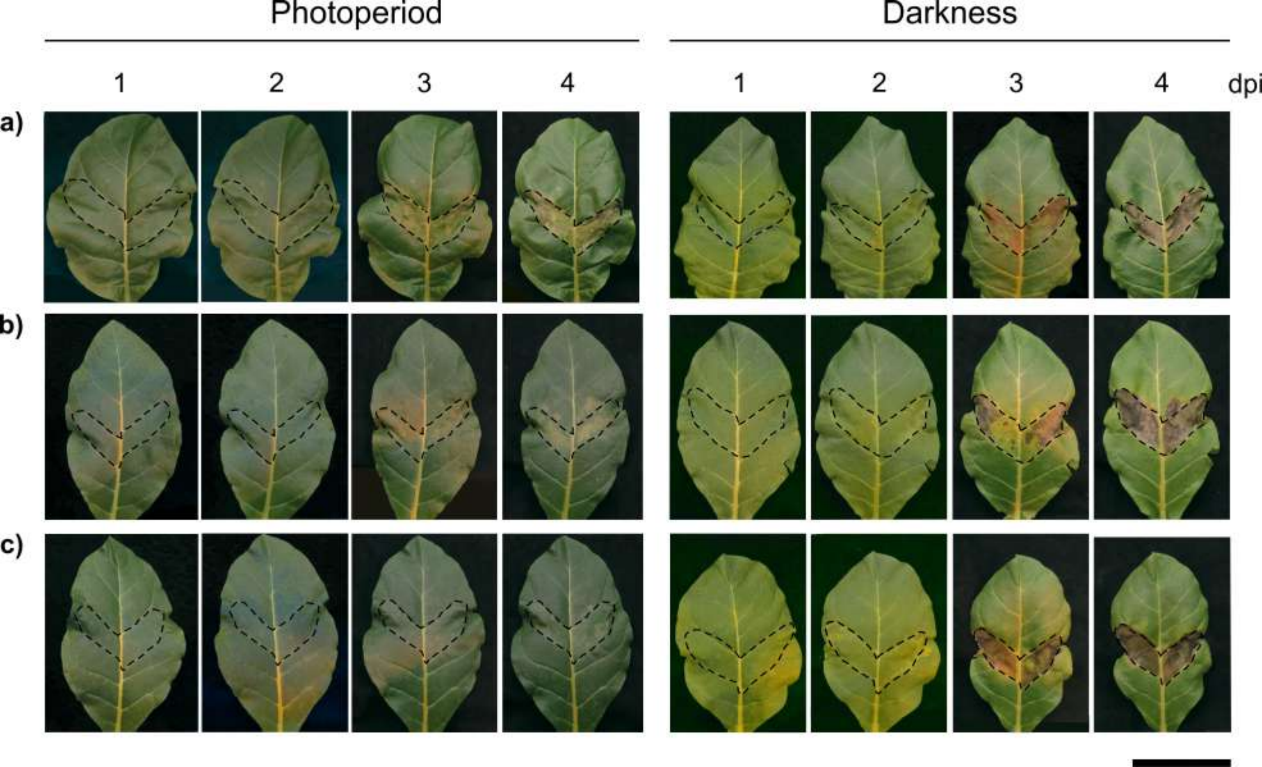

### Supplementary Figure S2

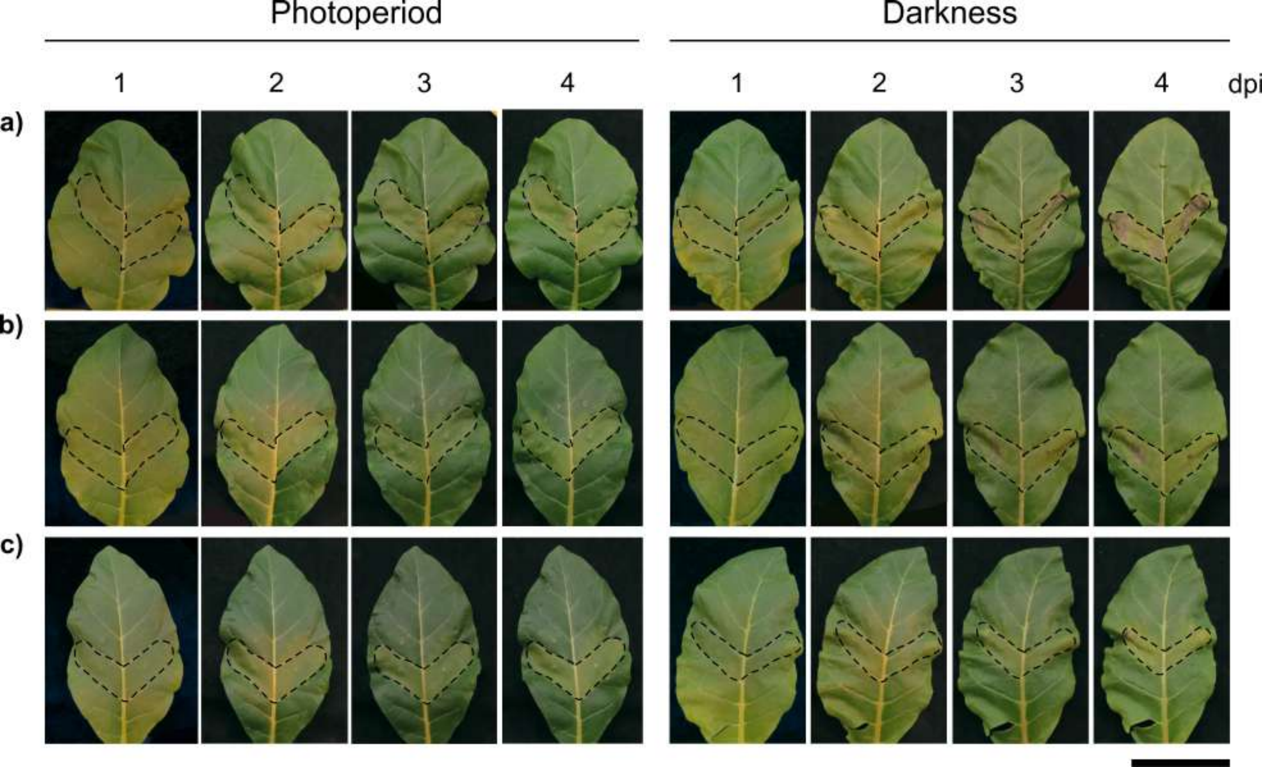

### Supplementary Figure S3

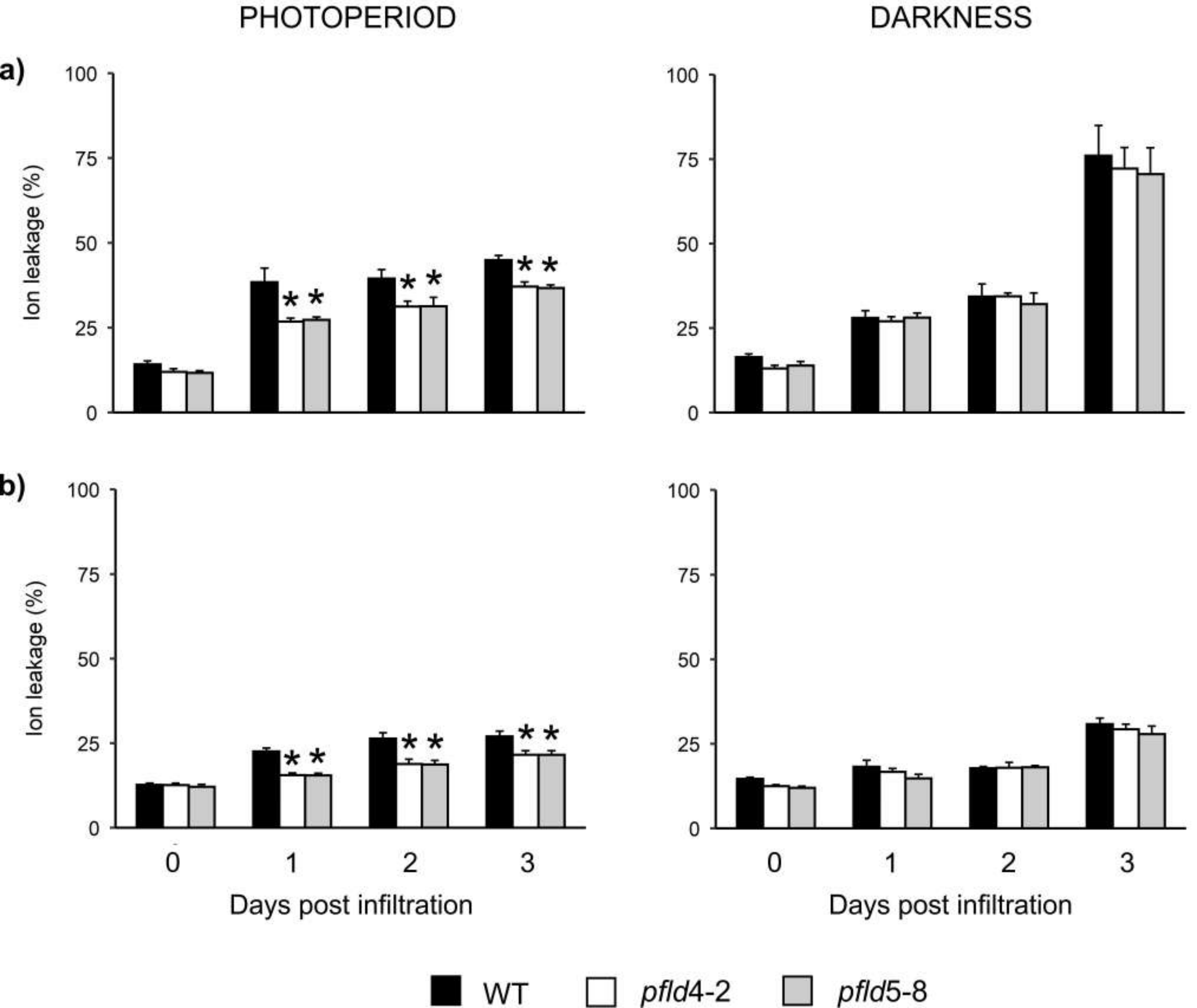

### Supplementary Figure S4

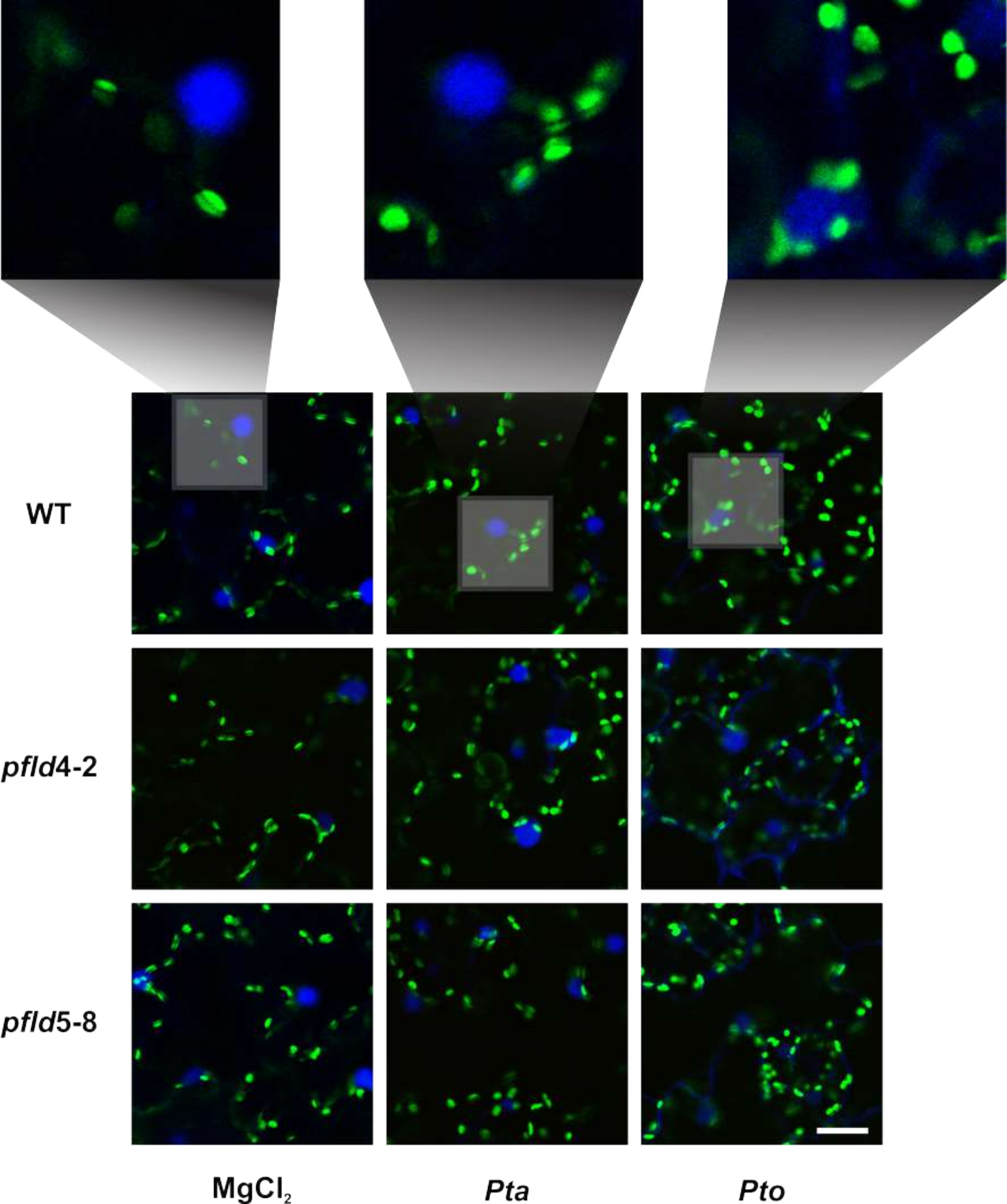

### Supplementary Figure S5

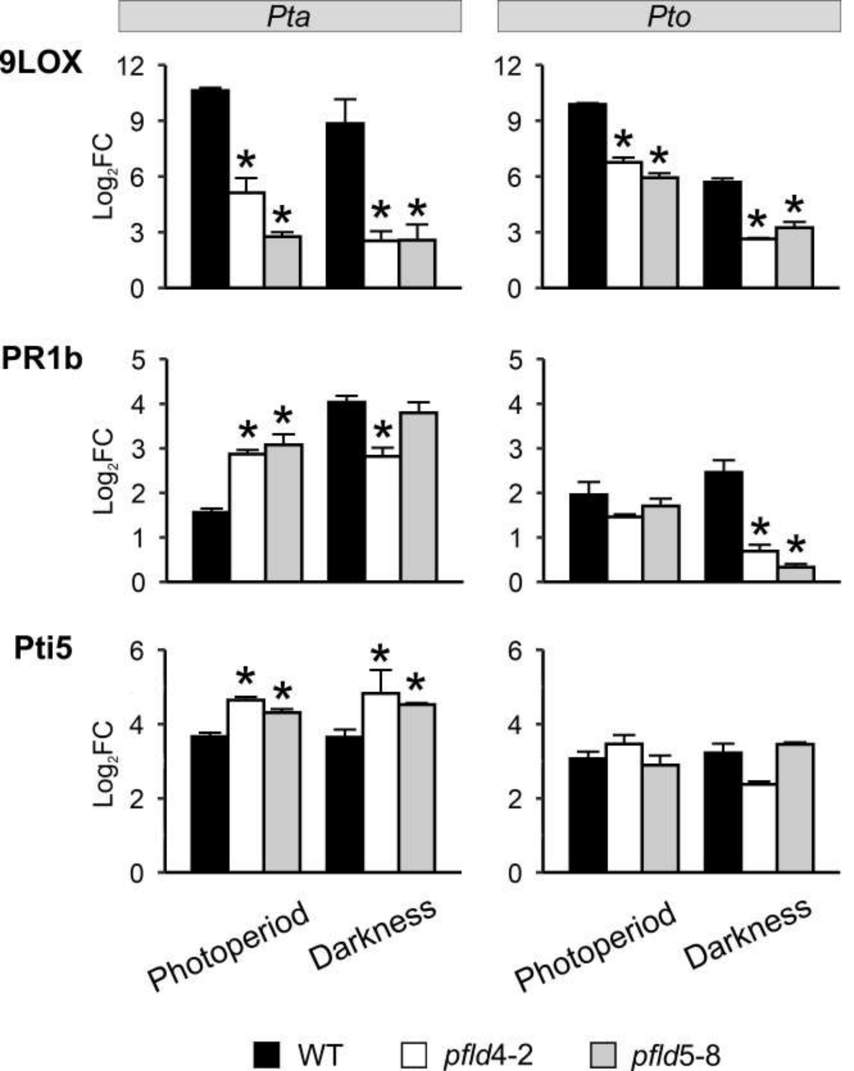

### Supplementary Figure S6

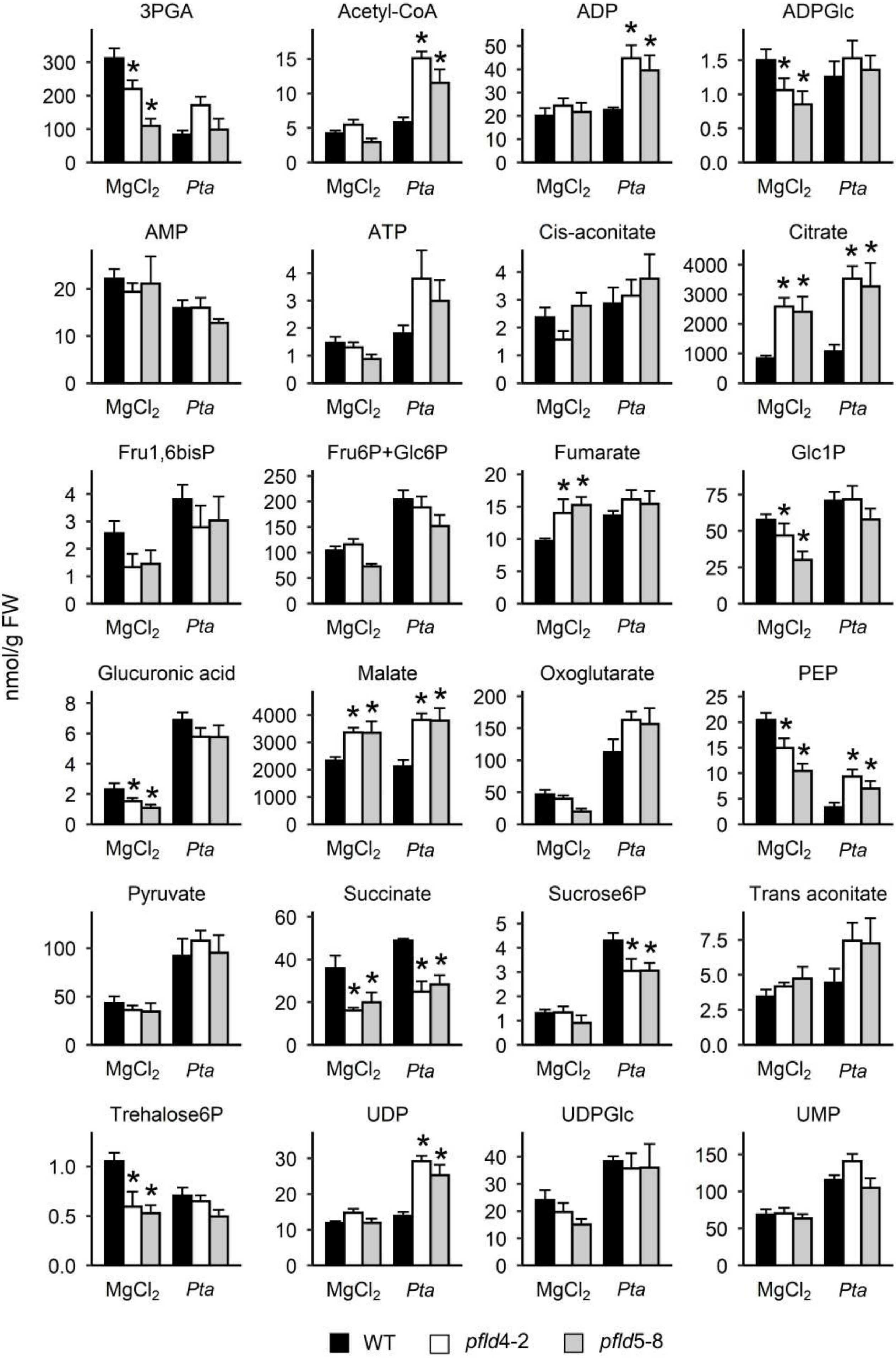

### Supplementary Figure S7

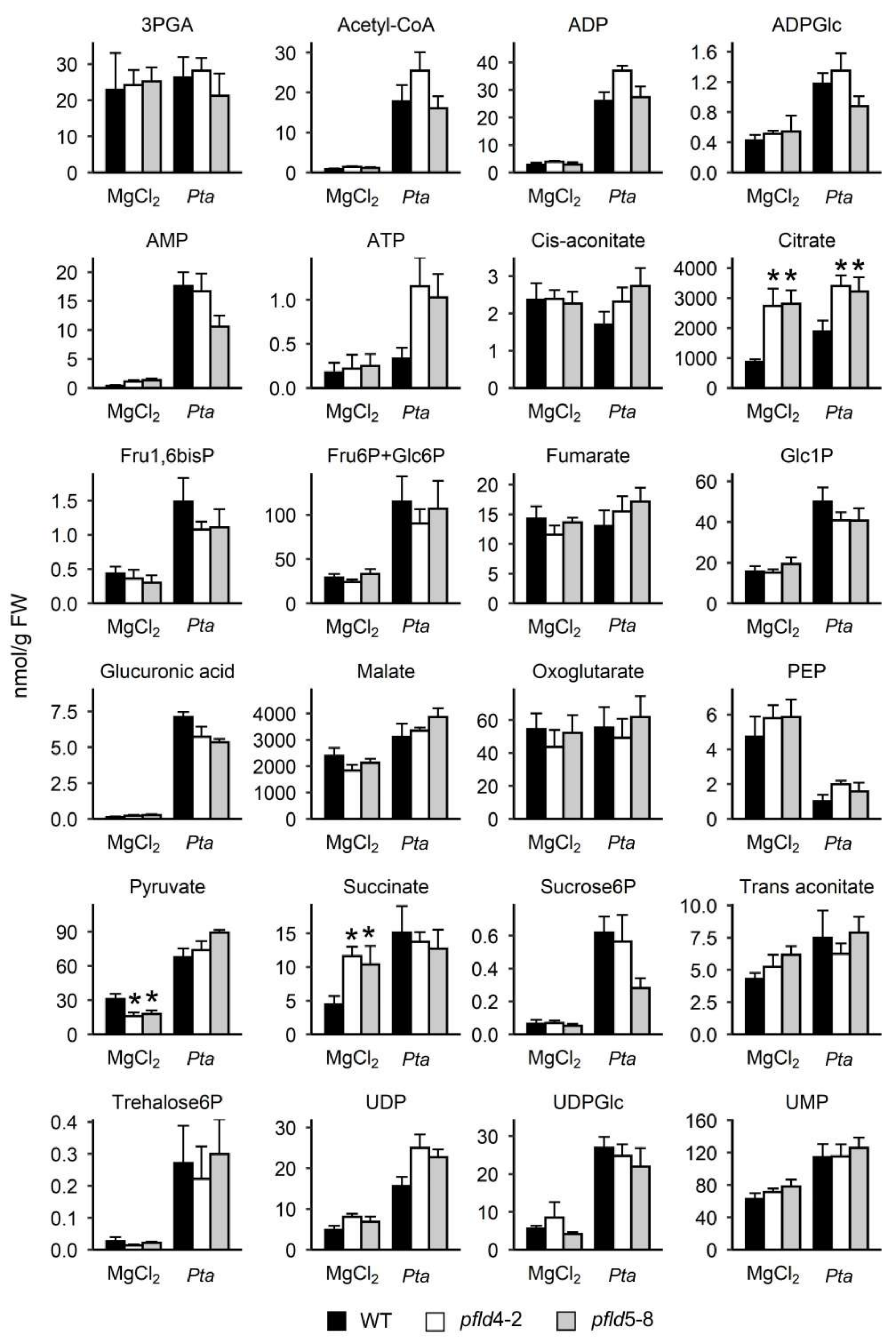

### Supplementary Figure S8

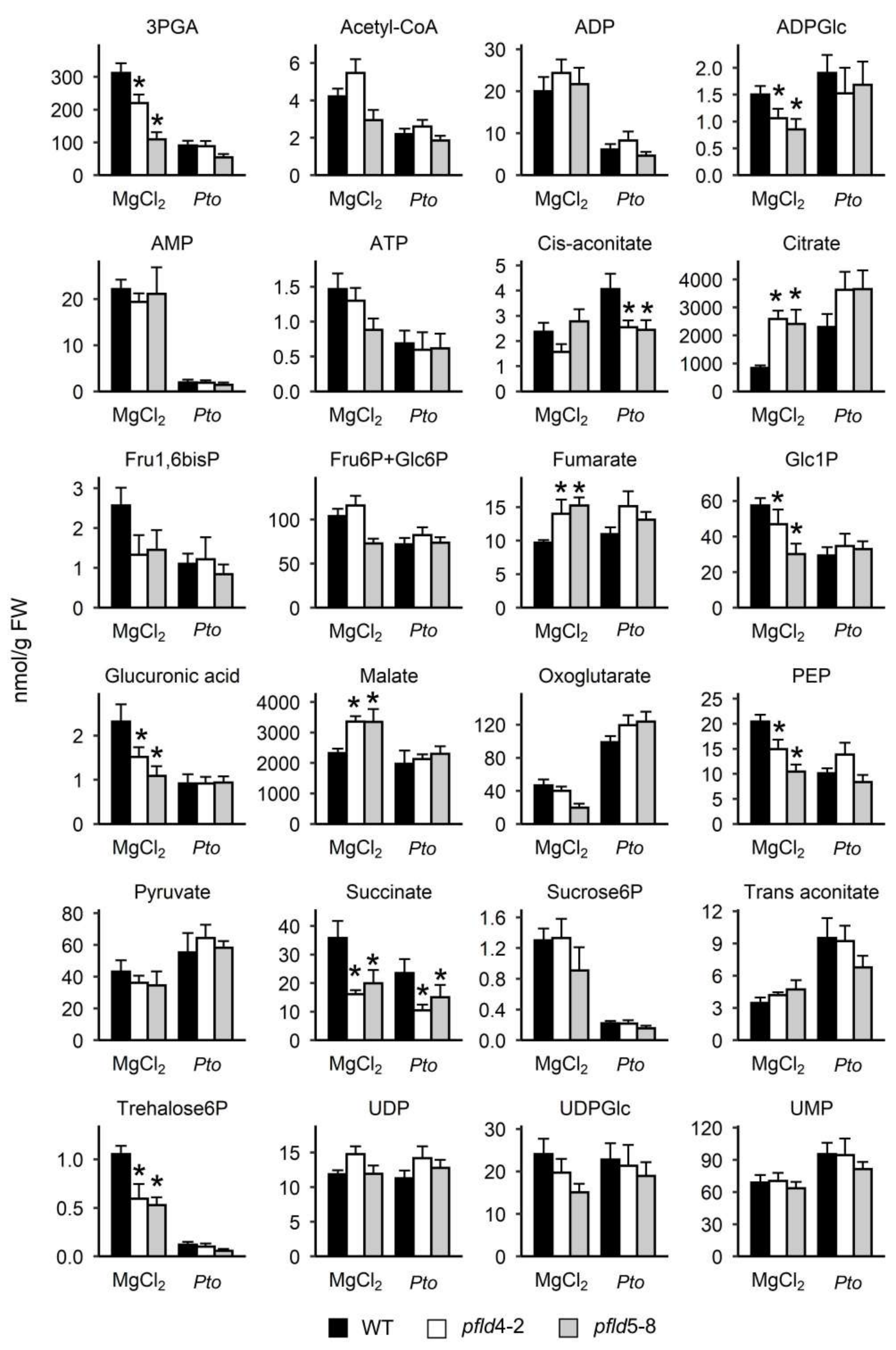

### Supplementary Figure S9

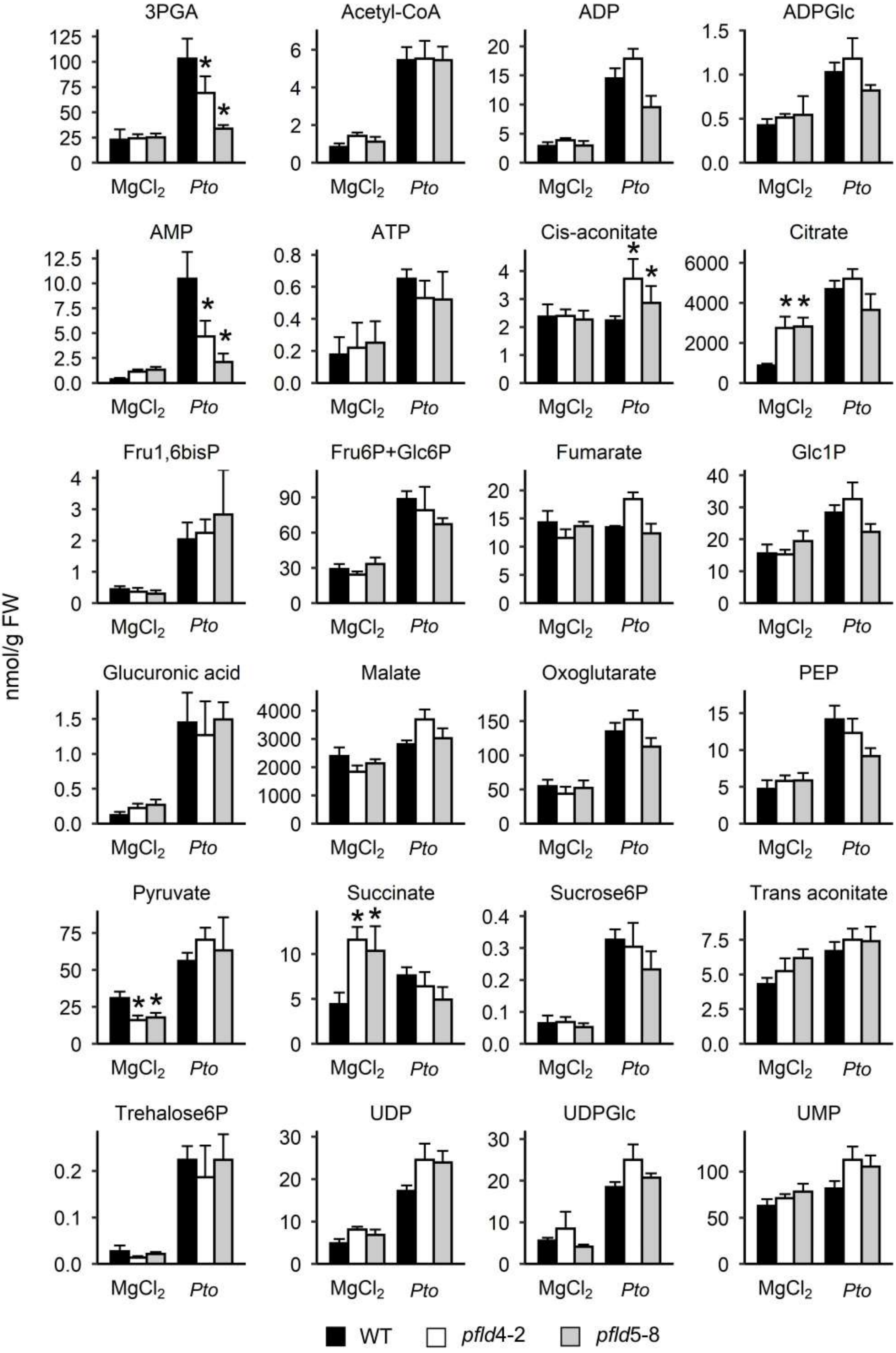

### Supplementary Figure S10

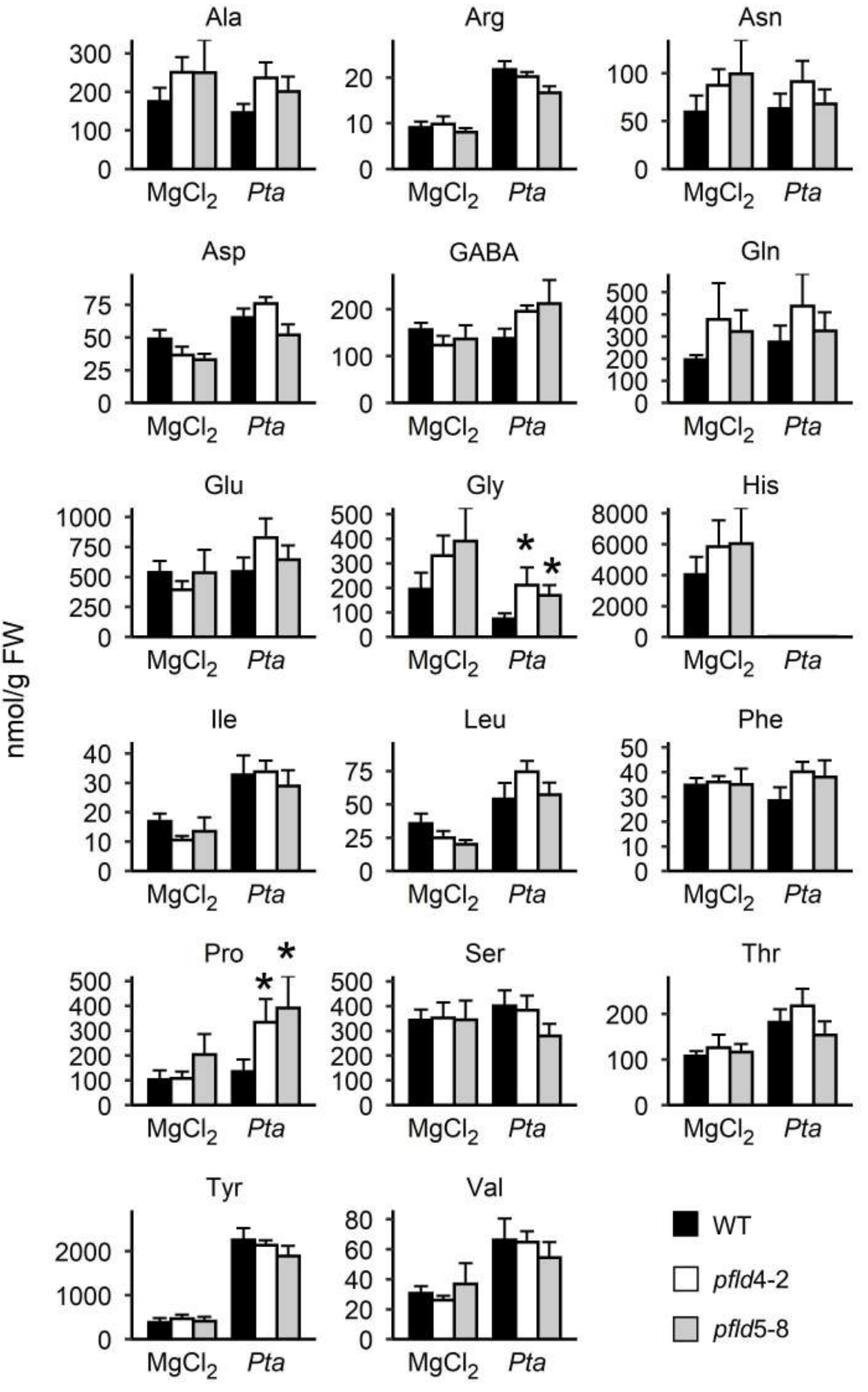

### Supplementary Figure S11

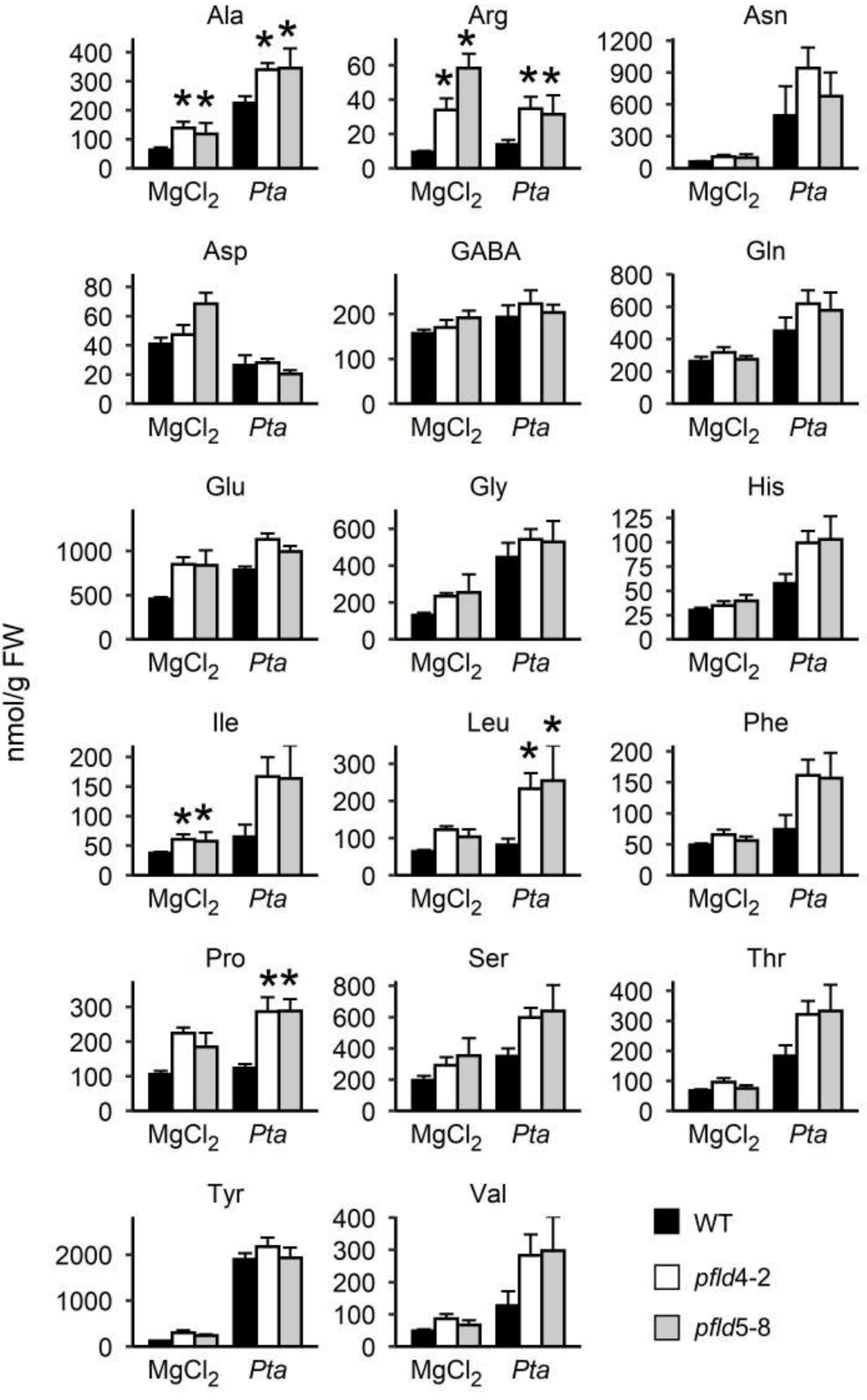

### Supplementary Figure S12

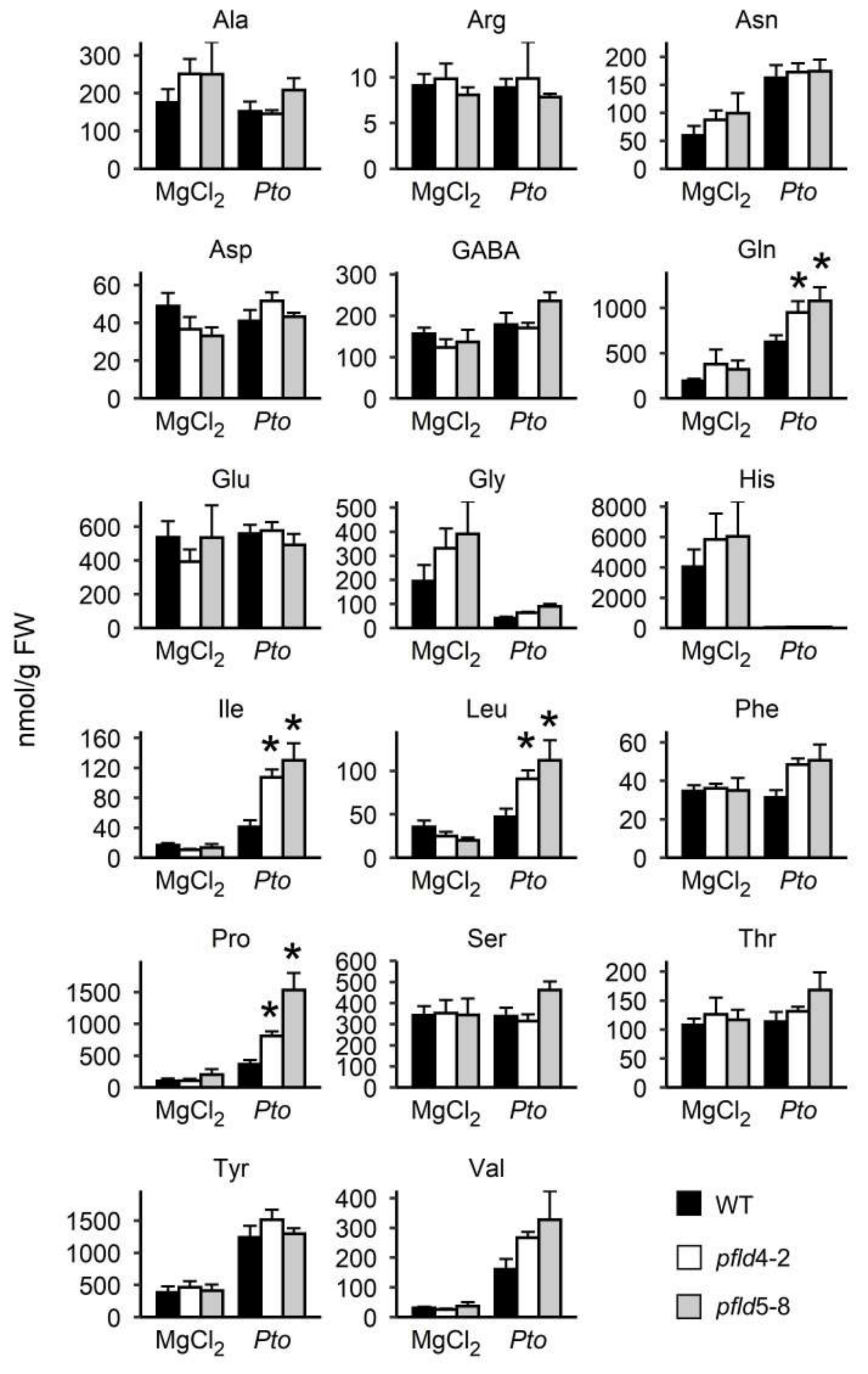

### Supplementary Figure S13

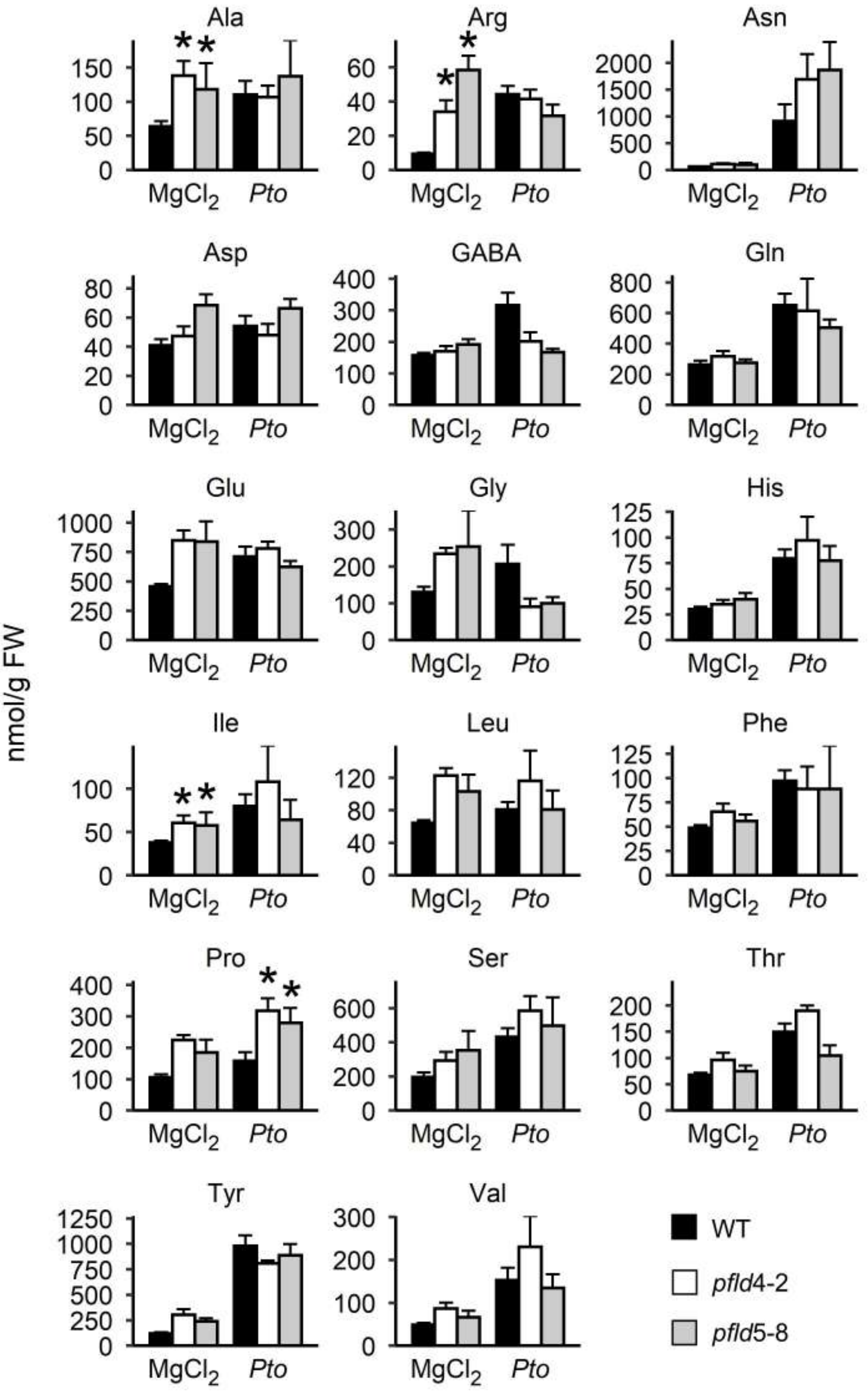
